# Daily ranging and den usage patterns structure fission-fusion dynamics and social associations in spotted hyenas

**DOI:** 10.1101/2021.10.01.462772

**Authors:** Eli D. Strauss, Frants H. Jensen, Andrew S. Gersick, Mara Thomas, Kay E. Holekamp, Ariana Strandburg-Peshkin

## Abstract

Environment structure often shapes social interactions. Spatial attractors that draw multiple individuals may play a particularly important role in dispersed groups, where individuals must first encounter one another to interact. We use GPS data recorded simultaneously from five spotted hyenas (*Crocuta crocuta*) within a single clan to investigate how communal dens and daily ranging patterns shape fission-fusion dynamics (subgroup splits and merges). We introduce a species-general framework for identifying and characterizing dyadic fission-fusion events and describe a taxonomy of ten possible configurations of these events. Applying this framework to the hyena data illuminates the spatiotemporal structure of social interactions within hyenas’ daily routines. The most common types of fission-fusion events involve close approaches between individuals, do not involve co-travel together, and occur at the communal den. Comparison to permutation-based reference models suggests that den usage structures broad-scale patterns of social encounters, but that other factors influence how those encounters unfold. We discuss the dual role of communal dens in hyenas as physical and social resources, and suggest that dens are an example of a general “social piggybacking” process whereby environmental attractors take on social importance as reliable places to encounter conspecifics, causing social and spatial processes to become fundamentally intertwined.

## INTRODUCTION

Environmental features can have a strong influence on social processes [1,2]. Spatial structure constrains social structure because spatiotemporal proximity is required for social interaction [3]. Features of the environment can also promote social behavior by increasing the frequency or effectiveness of interactions, or deter social interactions by decreasing or preventing contact between conspecifics and effectively segmenting the environment. Consequently, social behavior occurs non-randomly in space [1,4]. When particular features of the environment are attractive for multiple conspecifics, these spatial attractors can serve as catalysts for social encounters, thus influencing social structure [5–7]. The variety of spatial influences on social interactions, and the potential for environment structure to shape social structure more broadly, motivate a need to understand the extent to which observed patterns of association are driven by social preferences versus habitat preferences and spatial constraints [1,2,8,9].

The role of spatial structure in driving social structure is likely to be especially pronounced in groups with fission-fusion dynamics. In these systems, individuals associate in subgroups that change composition frequently. Social systems with a high degree of fission-fusion dynamics (henceforth, fission-fusion societies) are taxonomically diverse, occurring in many birds [10], fish [11], and mammals [12,13], including human societies [14]. Fundamental building blocks of social structure in these societies are fission-fusion events, or changes in subgroup composition. Because of the fluidity of these systems, individuals can control their social environment by deciding with whom to associate. As a result, many fission-fusion societies show high degrees of relational complexity [15] – that is, individuals form differentiated social relationships with their group-mates. However, the fluidity of these societies also presents challenges to individuals. Whereas social partners are close-at-hand in more cohesive groups, individuals residing in fission-fusion groups can less reliably find social partners, especially if they prefer socializing with particular group-mates. As a result, group sleeping sites, clumped food resources, or other spatial attractors are likely to be areas of frequent fission-fusion events, and thus are expected to have dramatic effects on social structure in these societies. However, currently little is known about how these spatial attractors influence spatial structure in fission-fusion societies.

Spotted hyenas (*Crocuta crocuta*) offer an ideal system in which to explore the influence of spatial attractors on social structure. Spotted hyenas (henceforth, hyenas) are carnivores living in closed groups that defend a common territory and show a high degree of fission-fusion dynamics [8,16]. Hyena groups are composed of multiple matrilines, with high relatedness within matrilines but low average relatedness within the clan [17]. Although clans can contain up to 126 individuals [18], hyenas associate in much smaller subgroups, and are often found alone [19]. Fission-fusion dynamics help individuals reduce the costs while retaining the benefits of sociality. Spending time alone or in small subgroups reduces feeding competition [19,20], and may also protect young offspring from infanticide [19,21]. However, hyenas form larger subgroups to engage in cooperative behavior such as interclan conflict, competition with sympatric carnivores, or patrolling territorial boundaries [19,22]. It is not only subgroup size that is relevant for spotted hyenas, but also which individuals are present in the subgroup. Spotted hyenas exhibit differentiated social relationships that are correlated with tolerance during feeding competition [23] and variation in social support, which is linked to dominance rank and fitness [24]. Therefore, the challenge of finding and interacting with social partners is critical in this species.

There are multiple, non-exclusive mechanisms by which hyenas might find and interact with preferred social partners. They could rely on chance encounters with other group members within their territory, then preferentially move together with preferred clan-mates. They could also coordinate non-chance meetings with specific social partners via long-distance communication facilitating convergence with those individuals [25], as in bonobos [26]. Finally, they could take advantage of spatial attractors, where the routine, predictable convergence of large subsets of the group facilitates locating and interacting with preferred partners. In support of the latter mechanism, large subgroups of hyenas are often found at communal dens, suggesting that dens may serve as spatial attractors that influence social structure. Communal dens are large complexes where females keep their young offspring until they are 10-12 months old [27,28]. Mothers with dependent offspring visit the den daily to nurse, and though hyenas do not engage in allocare, individuals without young offspring nevertheless visit the den regularly [28].

Although dens are known to be socially important for hyenas, the extent to which dens drive aggregate fission-fusion dynamics and social interaction patterns remains unknown. To address this question, we need (a) a system for monitoring the movements of multiple group members at once, and (b) an analytical approach for defining and quantifying fission-fusion events – the events by which subgroups of hyenas come together or split apart. Here we use multi-sensor collars to track the movements of multiple hyenas within the same clan simultaneously. We develop a general framework for modeling fission-fusion events between pairs of individuals as consisting of three canonical phases, distinguished by changes in distance between the two individuals over time. Using the movement characteristics of the two individuals in each phase, we construct a “taxonomy” of fission-fusion event types and compare qualitative and quantitative features of events that occur at communal dens to events that occur away from them. Next, using permutation-based reference models that preserve patterns of daily movement [29] and den usage, we explore the extent to which features of observed dyadic fission-fusion events can be explained by these factors. Finally, we compare social networks constructed from the fission-fusion events in the observed data and the reference models to understand the contribution of daily movement and den usage patterns to overall social structure.

## METHODS

### Data collection

We used custom-built tracking collars to collect data on the movements of five wild adult female spotted hyenas who were members of the same clan in the Masai Mara National Reserve, Kenya. These hyenas were observed as part of the Mara Hyena Project, a long-term study of several hyena clans ongoing since 1988. Individuals in this study were monitored near-daily from birth, providing important information about genealogical [30] and dominance relationships among members of the group [24]. To obtain a representative sample, we collared individuals who were not closely related or closely positioned in the dominance hierarchy. Daily monitoring data were used to identify communal dens, and hyenas in our study group were observed using four different communal dens over the course of the study (Figure 1C, Figure S1). Reproductive state varied among the study individuals -- two had den dependent cubs for the whole study, one gave birth halfway through the study, one was between reproductive events, and one had a den-independent but still nursing cub.

**Figure 1.**
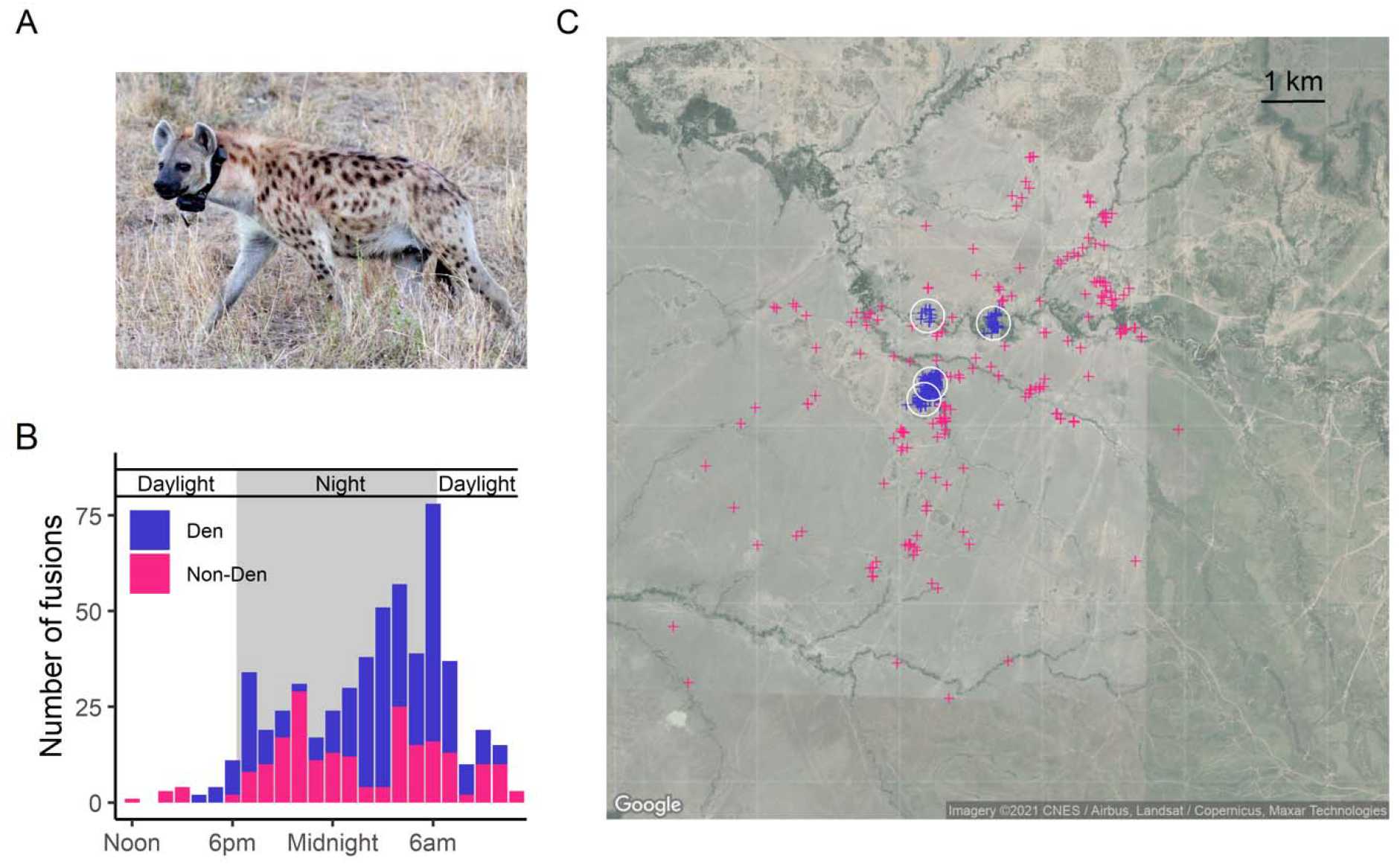
Spatial and temporal patterns of fission-fusion in spotted hyenas. (A) Female hyena wearing a tracking collar. (B) Time of day and (C) locations of the starts of fission-fusion events across all hyena pairs. Color specifies whether events started at a den (blue) or not (magenta). White circles represent locations of the four communal dens in use during the study period. See also Supplementary Video 1.

Each hyena wore a Tellus Medium collar (Followit Sweden AB) containing a custom-built sound and movement module modified from a DTAG board [31,32] and integrating a high-resolution (95% of points within <5m), high sample rate GPS (Gipsy-5 module, Technosmart, Italy), from which we use the GPS (1 Hz) and triaxial accelerometer (1000 Hz, downsampled to 25 Hz) data in this study. Collars were deployed throughout December 2016, and recorded continuously from January 1 until mid-February 2017 (Table S1, Supplementary Video 1). Prior to analysis, we performed minimal pre-processing of the GPS data to remove unrealistic locations and fill in very short gaps. The GPS data also contained some 12-hour gaps due to a firmware bug - such missing data accounted for 18% of the total tracking time. See Supplementary Material 1 and 2 for more details on collar specifications, collar deployment, data preprocessing, and missing data.

### Data analysis

We used the GPS data to identify fission-fusion events involving each pair of hyenas, and mapped the spatial and temporal distribution of these events, specifically in relation to dens. We then characterized the dynamics of these events in two ways. First, we developed a framework for breaking each event up into discrete phases, then categorized and combined these phases to produce a “taxonomy” of event types. Second, we quantified the properties of events via several continuous metrics (Table S2) and analyzed the distribution of these metrics for events occurring both at and away from dens. Next, we tested how well reference models that accounted for den usage and daily ranging patterns could capture the typical properties of events, as well as aggregate interaction patterns. Lastly, we constructed a social network based on the frequency of fission-fusion events for pair of individuals and compared this network to networks produced by our reference models.

#### Identifying fission-fusion events

Fission-fusion events are defined as sequences where two or more individuals come together (‘fusion’) for a period of time and later separate (‘fission’). Here, we extracted fission-fusion events at the dyadic level. For each pair of individuals, we identified contiguous periods of time where they came within a distance of 100m of one another. We identified the start of the event as the time when the distance between the individuals dropped below 200m (on its way down to < 100m) and the end as the time when this distance again rose above 200m. Using two thresholds avoided the problem of introducing many short “events” via individuals crossing a single threshold multiple times due to noise or small movements. We chose these thresholds to be consistent with definitions used for direct behavioral observations in the field, where individuals are considered together when within 200m of one another [33]. Changing these thresholds to 50m / 100m (inner / outer threshold) or 200m / 300m, while changing the total number of events and the exact ordering of event type frequencies, does not qualitatively change the overall observed patterns or interpretation (Supplementary Material 5).

#### Temporal and spatial distribution of fission-fusion events, and their relationship to dens

To characterize daily patterns in the occurrence of fission-fusion events, we counted the number of fusion events occurring across all dyads during each hour of the day.

To characterize where fission-fusion events typically occur, we classified fission-fusion events according to their spatial proximity to dens (Figure 1C), defining “den events” as events that either started or ended within 200m of a den, with the remaining events considered “non-den events”.

#### Analyzing the dynamical properties of fission-fusion events

The dynamics observed during dyadic fission-fusion events are complex and varied, yet all events share common features. The distance between the two individuals by definition follows a U-shaped structure during a fission-fusion event (Figure 2B), declining as they converge (*fusion phase*), remaining small while they spend time together (*together phase*), then rising again as they part ways (*fission phase*). We took advantage of this canonical structure to identify three phases for each event (fusion, together, and fission) by fitting a U-shaped function constructed of three line segments (Figure 2B) using constrained piecewise linear regression with least squares minimization (Supplementary Material 1). We constrained the three segments such that the first started at 200m, the last ended at 200m, and the middle segment had a slope of 0.

**Figure 2.**
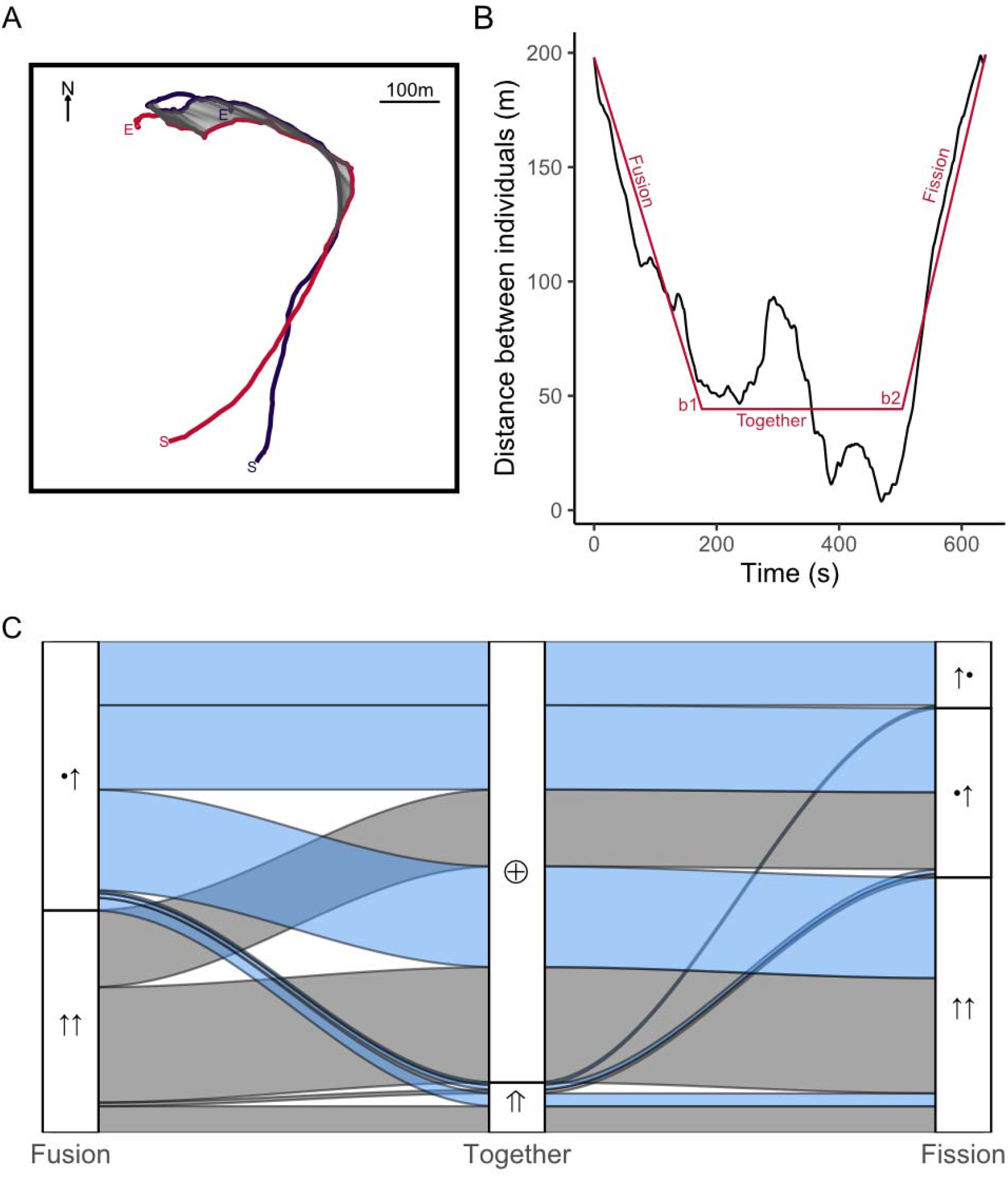
Example fission-fusion event and frequencies of event types. (A) Example trajectories of two individuals during an extracted fission-fusion event. Gray lines connect time-matched GPS points during the *together phase*. See Supplementary Videos for animated examples. (B) Distance between the two individuals over time (black) and fitted piecewise regression model (red). Break points are used to identify the three phases for each event. (C) Alluvial plot of the frequencies of transition motifs between different categories of the three phases. Symbols indicate the movement patterns of the two individuals involved in the event (• = stationary, ↑ = moving, ⊕ = local, ⇑ = traveling). Note that asymmetrical fissions can occur in two ways: either the two individuals show the same movement patterns as in the fusion phase (•↑), or the individuals reverse movement patterns (↑•).

As an output of this fitting procedure, we identified transition times (*b*_*1*_ and *b*_*2*_) which allowed us to decompose each event into three phases. Specifically, we defined the *fusion phase* as the time from the beginning of the event, *t*_*0*_, to *b*_*1*_; the *together phase* as the time from *b*_*1*_ to *b*_*2*_; and the *fission phase* as the time from *b*_*2*_ to the end of the event, *t*_*f*_.

After identifying phases, we used basic features of each phase to build a “taxonomy” of fission-fusion event types. As a basic descriptor of the *fusion* and *fission phases*, we identified which individual(s) moved during that phase. We classified the *fusion phase* into two *phase categories* - both individuals moved (↑↑), or one was stationary while the other moved (•↑). An individual was classified as having “moved” if its displacement between the beginning and end of the phase was greater than 5m, an upper bound on our estimated GPS error. The *fission phase* had three potential categories: both moving (↑↑), one stationary and one moving in the same arrangement as the *fusion phase* (•↑), or one moving and one stationary but with the movement roles reversed (↑•). Note that it was not possible for both individuals to remain stationary during these phases, as movement of at least one individual is necessary to result in a fusion or fission. We classified the *together phase* into “traveling” (⇑) if the individuals had a displacement of greater than 200 m during the phase, or “local” (⊕) if not.

Using these broad categories for each phase, we then analyzed the typical sequences of phase types seen in our data (Figure 2C).

Combining the categories of each of the phases allowed us to classify each event into one of ten distinct *event types* (Figure 3B). We can interpret these event types as broad classes of movement interactions between two individuals.

**Figure 3.**
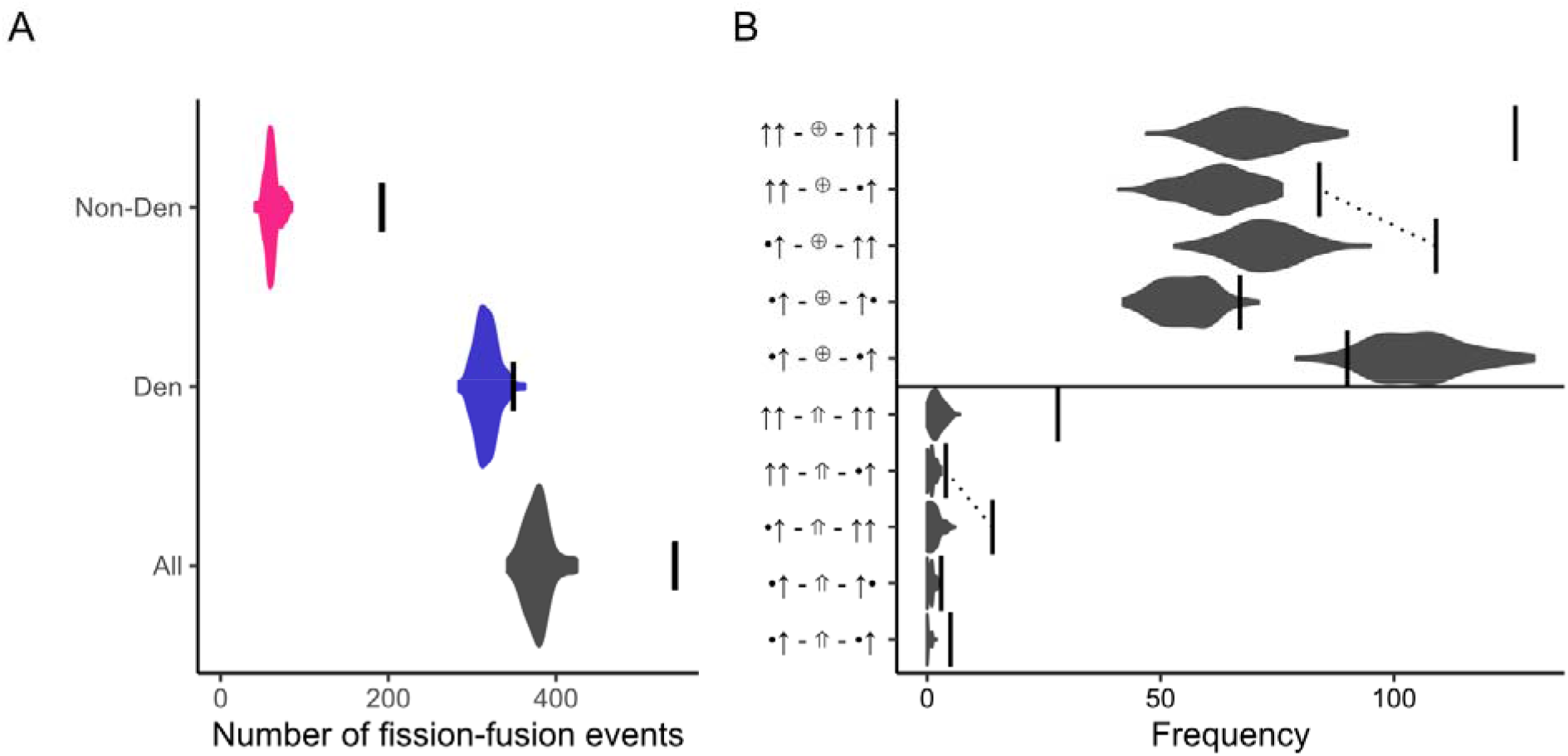
Comparison of the number of fission-fusion events in real data vs. reference models preserving den attendance and daily ranging patterns. (A) Overall number of events observed across real data (vertical lines) and reference models (violin plots) across all events, den events, and non-den events. (B) Frequency of events (x-axis) broken down by type (y-axis) in the real data (vertical lines) as compared to the reference models (violin plots). Y-axis labels represent the behavior of the two individuals during the three phases (from left to right: fusion, together, fission) for each event type. Dotted lines connect pairs of event types that are essentially time-reversed versions of each other, to highlight the asymmetry between fusions and fissions (see text).

For example, one possible event type is the category in which “one individual approaches another and then leaves” (Supplementary Video 2):

1. Fusion phase: one individual is stationary, one is moving: “•↑”
2. Together phase: local: “⊕”
3. Fission phase: the stationary individual remained stationary, the moving individual continued moving: “•↑”

We represent the complete event-type graphically as “•↑ - ⊕ - •↑.”

Alternatively, the event type •↑ - ⇑ - ↑↑ represents one individual approaching another that was stationary, then the two moving off and traveling together before mutually parting ways (Supplementary Video 3). After classifying events into types, we analyzed how often each event type occurred in our data to assess what types of fission-fusion events are characteristic of hyena interactions.

We also characterized the properties of events through continuous metrics: event duration; displacement, directional synchrony and activity synchrony during the *together phase*; and distance from the den at the start and end of the event (Table S2).

#### Constructing association networks based on fission-fusion events

To quantify broader patterns of social structure amongst our tracked hyenas, we constructed a social network. We defined edge weights using a version of the simple ratio index [34]:

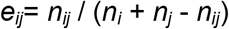

where *n*_*ij*_ represents the number of fission-fusion events involving individuals *i* and *j, n*_*i*_ the number of events involving individual *i* and any other individual, and likewise for *n*_*j*_. This metric quantifies the extent to which *i* and *j* associate with each other as a fraction of their associations with all other tracked individuals.

#### Permutation-based reference models for fission-fusion events

To test to what extent den usage and daily ranging patterns underlie the observed properties of fission-fusion events at the individual-event and aggregate levels, we constructed permutation-based reference models. To do so, we permuted our data such that the trajectory of each individual for a given day was randomly assigned to a different day [29]. This permutation preserves each individual’s overall ranging patterns and typical daily patterns of movement, but breaks the temporal link between the trajectories of pairs of individuals. Because communal dens changed partway through the study, we accounted for den usage by constraining the permutation to only swap days from periods where the individual was using the same den or set of dens (Figure S1). We also constrained the permutations such that no two individuals were randomly “matched” to the same day, thus ensuring a complete break-up of the temporal links between trajectories. To minimize possible artefacts arising from temporal discontinuities at the “break point” between days, we used noon as the break point (because hyenas are generally least active around mid-day) and also removed any events crossing noon in both observed data and reference models from all analyses involving the reference models.

For each reference model (n = 100 instantiations), we carried out the same analyses as described above (i.e. extracting fission-fusion events, characterizing their phases and types, computing their properties, and constructing a social network) to allow comparison with the real data.

## RESULTS

### When and where do fission-fusion events occur?

Overall, we identified 690 fission-fusion events across the five tracked hyenas in our study, for 551 of which we could identify the exact start and end times enabling further analysis (Supplementary Material 1). Fission-fusion events were more likely to occur at night than during the day, with peak occurrence around dusk and dawn (Figure 1B). Median duration of the *together phase* of fission-fusion events was 20.58 (IQR = 3.23 - 68.48) minutes. Analyzing the spatial distribution of these events (Figure 1C) revealed that 62% of fusions (n = 339 events) and 57% of fissions (n = 315 events) occurred at a communal den, with a total of 64% of all events either starting or ending at a den (n = 350 events).

### What types of fission-fusion events are observed?

Our phase categorization scheme (Figure 2C) revealed that for most events (89%), the *together phase* was local (⊕), i.e. the two individuals did not travel more than 200m while together. For events where the together phase involved travel (⇑), these were approximately equally likely to be initiated by one individual moving to meet another (•↑) as by both initially moving and their paths converging (↑↑). However, these traveling events most often ended with both individuals continuing moving and their paths diverging (↑↑), rather than with a single individual remaining stationary while the other moved off (•↑ or ↑•). Of events involving joint travel, 36% started at the den and 9% ended at the den, indicating that individuals more often met up at the den and traveled elsewhere than traveled together to the den.

Categorizing full events into types revealed clear differences in the relative frequencies of different fission-fusion configurations (Figure 3B). Although the definition of fission-fusion events is symmetric (one could think of a fission as simply a fusion in reverse), our data revealed an asymmetry in how fissions vs. fusions occur (Figure 3B; compare events connected by dashed lines). It was much more common for individuals to engage in an interaction where one was initially stationary in the *fusion phase* and later both moved off during the *fission phase* than the reverse. In other words, it was much more common for individuals to meet by arriving in sequence to a given location and then move off at the same time than it was for them to arrive synchronously and leave asynchronously.

### To what extent are fission-fusion patterns explained by individual daily ranging and den usage?

Our permutation-based reference models revealed that a large fraction of the observed number of fission-fusion events would be expected purely based on daily ranging and den usage patterns (Figure 3A). In particular, the reference models predicted a median of 379 events (95% range: 349 - 418), whereas the real data contained 543 events (70%). When considering only events starting or ending at dens, the reference models predicted a median of 316 events (95% range: 295 - 345), compared to 350 den events in the real data. Thus, the reference models accounted for approximately 90% of den events. In contrast, the reference models predicted a median of 60 (95% range: 47-80) events occurring away from dens, capturing only 31% of them compared to the real data (193 non-den events).

There was clear variation in how well the reference models captured different types of events (Figure 3B). While most event types were underrepresented in the reference models compared to the real data, the number of local events where one of the individuals remained stationary during both the fusion and the fission phase (•↑ - ⊕ - •↑, Supplementary Video 2) was actually slightly overrepresented. Conversely, events involving both individuals moving off during the fission phase were particularly underrepresented. Despite the overall lower number of events, the relative frequency of different event types observed in the data was roughly captured by the reference models.

When quantifying continuous properties of fission-fusion events (Table S2), the reference models captured some properties much better than others (Figure 4). Specifically, the distribution of event durations was approximately the same in the reference models as in the real data (Figure 4A), as was the distribution of events across the day (Figure 4D). However, displacements of individuals during the together phase were in general greater during real events than during artificial events generated by the reference models (Figure 4B), and hyenas approached each other much more closely in the real data (Figure 4C). Hyenas also had higher heading similarity (Figure 4E) and activity synchrony (Figure 4F) during real events, although activity synchrony at the den was reasonably well captured by the reference models.

**Figure 4.**
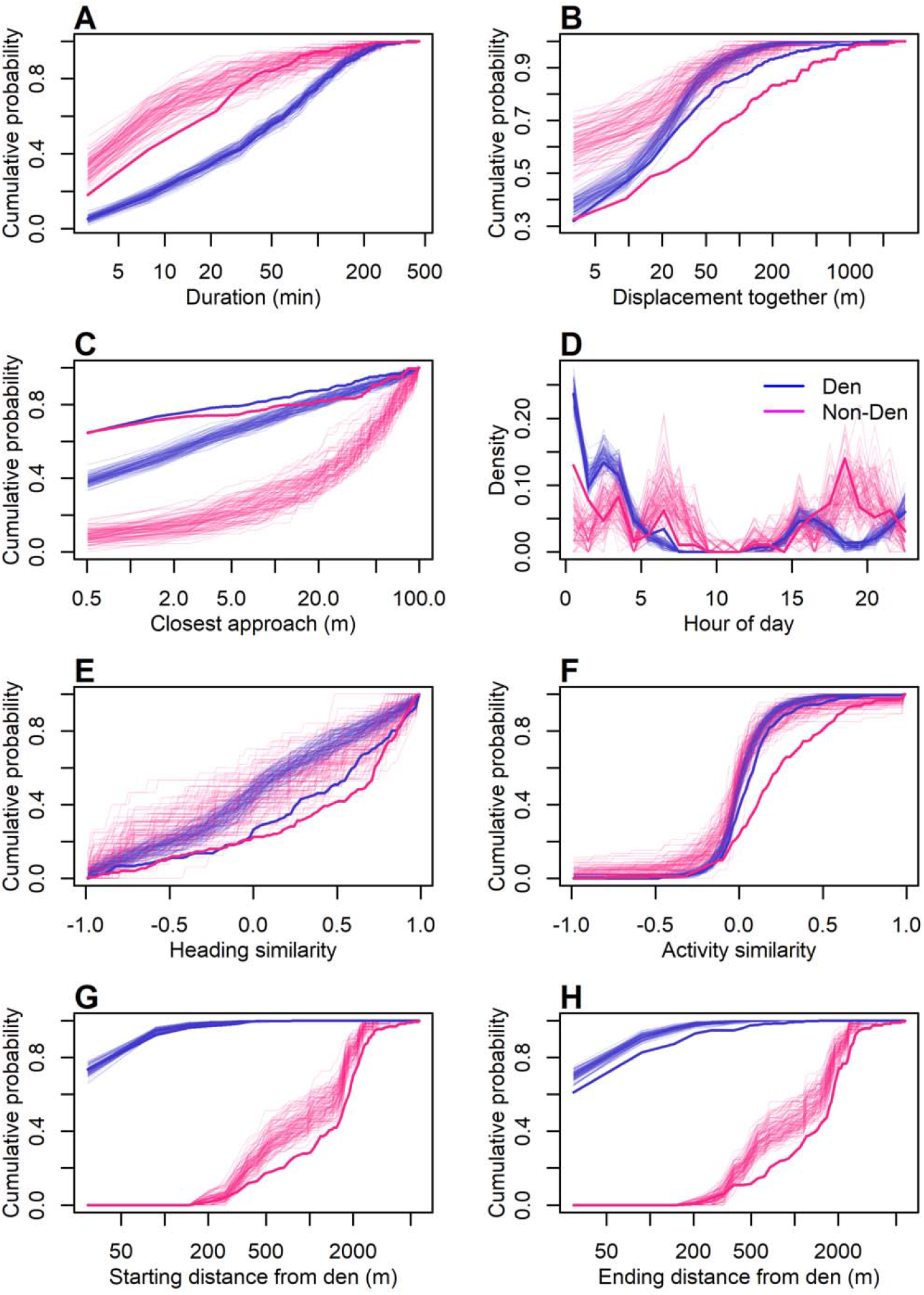
Reference models reproduce some but not all properties of observed fission-fusion events. Comparison of the detailed properties of fission-fusion events in the real data (thick lines) and in reference models (thin lines), broken up by whether the event occurred in the vicinity of a den (blue) or not (magenta). Plots show the cumulative distribution of each metric (x-axis labels) across all events. Note that panel D shows the distribution rather than the cumulative distribution.

Scaling up from individual events to social networks, we found that the reference models accurately captured the overall patterns of association amongst the individuals in our study (Figure 5). Empirically measured edge weights in an association network all fell within the expected ranges of values from the reference models, and the overall relative ranking of edge weights across dyads in the reference models was consistent with that observed in the real data. These results indicate that accounting for daily ranging and den usage patterns alone was enough to broadly explain the association patterns of the spotted hyenas observed in this study.

**Figure 5.**
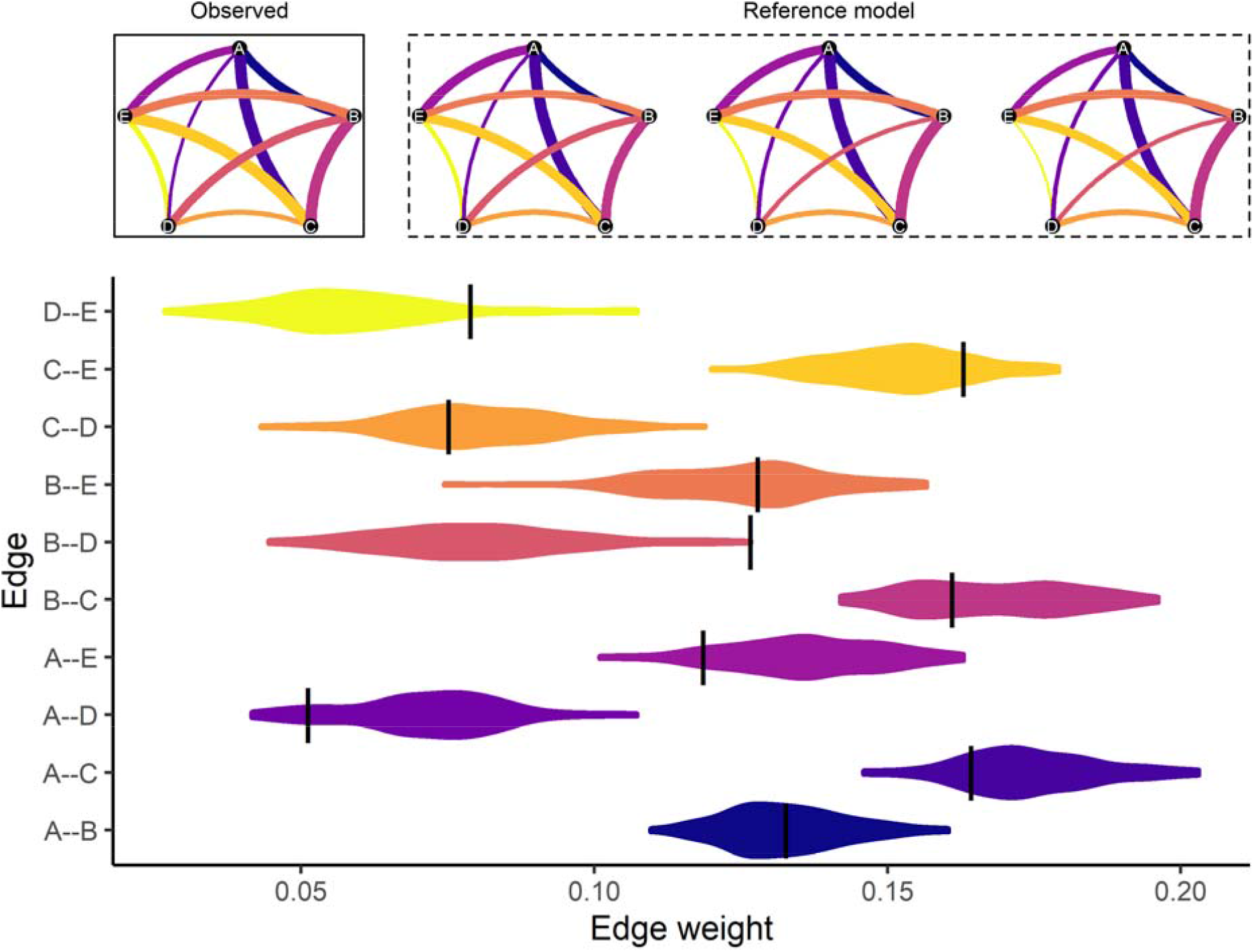
Reference models reproduce differentiated relationships found in observed social networks built from fission-fusion events. Black lines indicate observed edge weight representing frequency of association. Violins depict distributions of edge weights in 100 instances of the reference model.

## DISCUSSION

We present here a framework for measuring dyadic fission-fusion events and use it to quantify the extent to which individual daily ranging patterns, particularly those involving the communal den, underlie social structure in spotted hyenas. We found that over 60% of fission-fusion events either began or ended at the den, emphasizing that the communal den serves a critical role as a social hub for hyenas. We also found that reference models preserving daily movement patterns and den attendance were sufficient to explain many, but not all, features of the observed patterns of fission-fusion dynamics in hyena societies. Reference models captured the duration of events and the time of day when they occurred, but underestimated the overall frequency of events, the amount of synchrony between the individuals during the event, the closest approach between the individuals, and the occurrence of joint travel. Dyadic association strengths in the observed data were very similar to dyadic association strengths in networks created from the reference model, suggesting that social structure is closely linked to spatial structure in this species.

Overall, our results suggest that daily movement and den usage patterns strongly structure patterns of social encounters in hyenas. For a given hyena, it appears that which individuals it encounters and how long those encounters last follow in large part from its daily ranging patterns and visits to the communal den. However, the reference models underestimate the frequency of fission-fusion events, especially those not occurring near dens, suggesting that hyenas are more gregarious than expected based simply on spatiotemporal overlap. These models also underestimate the coordination of individuals during events, indicating that factors beyond spatial attractors – most likely social processes – underlie variation in what happens after hyenas encounter one another. Our findings suggest that hyenas may pursue a mixed strategy for acquiring critical social-interaction time with important partners, leaning heavily on the passive mechanism of co-occurrence with groupmates at the communal dens, but also actively pursuing convergence with particular individuals during the more dispersed, travel-heavy phases of the daily routine. This has important implications for hyena social structure, particularly in cases where a single group of hyenas has multiple active communal dens, as sometimes occurs (Figure S1). Our results suggest that the use of multiple communal dens should drive a more modular social structure. In fact, each of the four permanent group fission events documented in our study population since 1988 (e.g., [35]) was preceded by use of multiple communal dens, suggesting that this enhanced modularity may have important consequences for the fates of entire hyena societies.

Our results highlight the need for future work aimed at understanding the interplay between social and spatial structure. In particular, a shortcoming of this study is that we have data on only a few individuals. In order to achieve a broad sample of dominance ranks, we tagged individuals evenly distributed throughout the social hierarchy. However, because rank is closely associated with kinship in hyena societies, and kin tend to form the strongest social bonds [33,36], our study individuals were not closely bonded, and it remains unclear to what extent our results will generalize to more closely bonded individuals. A productive next step would be to deploy tags on many more individuals concurrently, ideally on every member of a social group. Doing so would also allow for the expansion of the approach presented here from dyadic fission-fusion events to polyadic events. Future work should also examine the role of more transient spatial attractors in contributing to social structure. For example, ungulate kills are ephemeral physical resources that lead to the aggregation of many group-mates, and are thus likely to be a large driver of fission-fusion events in hyenas. In addition, although fission-fusion events involving joint travel were uncommon, these events nonetheless may represent particularly important social events (e.g. group hunts or collective border patrols) or may be more common among strongly bonded individuals, and thus merit further study. For example, co-traveling may play an important role in the vertical transmission of social relationships from mother to offspring [37]. Finally, hyenas are known to use long-distance vocalizations to recruit their clan-mates over large distances in contexts requiring collective action [25], so the role of communication in driving fission-fusion patterns also warrants further investigation.

Our results provide important methodological insight into how to study social behavior in fission-fusion systems. We suggest that a useful approach is to distinguish drivers of social encounters from drivers of social interaction -- that is, an explicit distinction between the processes that drive (1) when, where, and which conspecifics individuals encounter (2) what individuals do and how long they spend together and (3) how and why they part ways. Our three-phase fission-fusion event model provides a useful tool for asking these questions by offering a means to identify and measure the fusion, together, and fission components of fission-fusion events. Furthermore, the taxonomy of event types derived from these phases facilitates understanding of processes operating across phases. For instance, our analysis of the frequency of different event types revealed a fundamental asymmetry between fissions and fusions (Figure 3B), indicating that fissions are not simply fusions in reverse. This approach provides a generalizable framework for future work investigating the dynamics of fission-fusion events across different social groups, species, and spatial scales.

Finally, the dual role of communal dens as physical and social resources suggests a potentially broadly-acting process by which spatial and social heterogeneity become aligned. By attracting individuals or promoting social interactions, spatiotemporally predictable resource hotspots become social hotspots, leading individuals to use these locations for social purposes. Through this “social piggybacking” effect, the social landscape conforms to the physical landscape, and socially-driven and resource-driven movements produce the same behavior. The communal den in spotted hyenas is a clear example of this process: the physical resource is only useful to a subset of individuals (mothers with den-dependent offspring), yet non-reproductive individuals frequently visit, demonstrating that this physical resource has become a social resource. When the resource in question is useful to all individuals, it becomes difficult to infer whether movements to it result from social or spatial processes, but both are likely to occur. For instance, foraging glades in vulturine guineafowl are sources of physical resources (food), but also serve as a hotspot of social interactions among groups, providing an opportunity for the movement of individuals or information across group boundaries [38]. Foraging sites, watering holes, resting sites, sunny/shady locations, or locations with good visibility for vigilance are all examples of spatiotemporally predictable physical resources that can become social resources. This dual socio-spatial process represents a challenge for existing paradigms for understanding drivers of animal associations. If social piggybacking occurs, null models such as the one we used here, which are typically interpreted as capturing spatial processes underlying animal movements, may also be capturing social processes. This suggests that in some contexts, it may be most meaningful to understand animal associations as a product of joint interaction between social and spatial processes rather than trying to disentangle them.

## ACKNOWLEDGEMENTS

We thank the Kenya Wildlife Service, the Narok County Government, and the Kenyan National Committee on Science, Technology and Innovation, the Naboisho Conservancy, the Mara Conservancy and Brian Heath for permissions to conduct this research. Thanks to many current and former members of the Mara Hyena Project for detailed data collection. We thank Benson Pion, Rebecca LaFleur, and Morgan Lucot for assistance in the field during collar deployment. We thank Mark P. Johnson for providing the DTAG tag technology and contributing to collar development and testing as well as discussions about ranging patterns and den use. We also thank the members of the Communication and Coordination Across Scales team for feedback and useful discussion.

## ETHICS

All field methods were approved by Kenya Wildlife Service under permit KWS/BRM/5001 to KEH. They conform to guidelines published by the American Society of Mammalogists [39], and they were also approved by the IACUC at Michigan State University under approval PROTO201900126, which was issued most recently in January, 2020.

## DATA, CODE, AND MATERIALS

Raw movement and accelerometer data used in this study are publicly available from Dryad data repository [40]. All analysis code required to replicate the results is available at https://github.com/arianasp/hyena_fission_fusion.

## AUTHOR CONTRIBUTIOI NS

ASG, FHJ, ASP, and KEH conceived the hyena collaring study, secured funding, and collected the tracking data that forms the basis for this work. KEH maintains the long-term study of spotted hyenas that enabled this work, and worked with FHJ to integrate and test the tracking collars. EDS and ASP conceived the specific questions addressed by this work, designed the analyses, wrote the analysis code, and produced a first draft of the manuscript. MT conducted a thorough code review to ensure the accuracy and reproducibility of analyses and implemented improvements to the analysis. All authors contributed to, and approved, the final manuscript.

## REFERENCES

1. He P, Maldonado-Chaparro AA, Farine DR. 2019 The role of habitat configuration in shaping social structure: a gap in studies of animal social complexity. Behav. Ecol. Sociobiol. 73, 9. (doi:10.1007/s00265-018-2602-7)

2. Webber QMR, Vander Wal E. 2018 An evolutionary framework outlining the integration of individual social and spatial ecology. J. Anim. Ecol. 87, 113–127. (doi:10.1111/1365-2656.12773)

3. Farine DR. 2015 Proximity as a proxy for interactions: Issues of scale in social network analysis. Anim. Behav. 104, e1–e5. (doi:10.1016/j.anbehav.2014.11.019)

4. Wolf JBW, Trillmich F. 2007 Beyond habitat requirements: Individual fine-scale site fidelity in a colony of the Galapagos sea lion (Zalophus wollebaeki) creates conditions for social structuring. Oecologia 152, 553–567. (doi:10.1007/s00442-007-0665-7)

5. Farine DR et al. 2015 The role of social and ecological processes in structuring animal populations: A case study from automated tracking of wild birds. R. Soc. Open Sci. 2. (doi:10.1098/rsos.150057)

6. Firth JA, Sheldon BC. 2015 Experimental manipulation of avian social structure reveals segregation is carried over across contexts. Proc. R. Soc. B Biol. Sci. 282, 20142350. (doi:10.1098/rspb.2014.2350)

7. Mourier J, Vercelloni J, Planes S. 2012 Evidence of social communities in a spatially structured network of a free-ranging shark species. Anim. Behav. 83, 389–401. (doi:10.1016/j.anbehav.2011.11.008)

8. Aureli F et al. 2008 Fission-Fusion Dynamics: New Research Frameworks. Curr. Anthropol. 49, 627–654.

9. Croft DP, Darden SK, Wey TW. 2016 Current directions in animal social networks. Curr. Opin. Behav. Sci. 12, 52–58. (doi:10.1016/j.cobeha.2016.09.001)

10. Silk MJ, Croft DP, Tregenza T, Bearhop S. 2014 The importance of fission-fusion social group dynamics in birds. Ibis (Lond. 1859). 156, 701–715. (doi:10.1111/ibi.12191)

11. Kelley JL, Morrell LJ, Inskip C, Krause J, Croft DP. 2011 Predation Risk Shapes Social Networks in Fission-Fusion Populations. PLoS One 6, e24280. (doi:10.1371/journal.pone.0024280)

12. Archie EA, Moss CJ, Alberts SC. 2006 The ties that bind: Genetic relatedness predicts the fission and fusion of social groups in wild African elephants. Proc. R. Soc. B Biol. Sci. 273, 513–522. (doi:10.1098/rspb.2005.3361)

13. Mann J, Connor RC, Tyack PL, Whitehead H, editors. 2000 Cetacean societies: field studies of dolphins and whales. University of Chicago Press.

14. Marlowe FW. 2005 Hunter-gatherers and human evolution. Evol. Anthropol. 14, 54–67. (doi:10.1002/evan.20046)

15. Lukas D, Clutton-Brock TH. 2018 Social complexity and kinship in animal societies. Ecol. Lett. 21, 1129–1134. (doi:10.1111/ele.13079)

16. Kruuk H. 1972 The Spotted Hyena: A Study of Predation and Social Behavior. Chicago, Illinois: The University of Chicago Press.

17. Van Horn RC, Engh AL, Scribner KT, Funk SM, Holekamp KE. 2004 Behavioural structuring of relatedness in the spotted hyena (Crocuta crocuta) suggests direct fitness benefits of clan-level cooperation. Mol. Ecol. 13, 449–458.

18. Green DS, Johnson-Ulrich L, Couraud HE, Holekamp KE. 2018 Anthropogenic disturbance induces opposing population trends in spotted hyenas and African lions. Biodivers. Conserv. 27, 871–889. (doi:10.1007/s10531-017-1469-7)

19. Smith JE, Kolowski JM, Graham KE, Dawes SE, Holekamp KE. 2008 Social and ecological determinants of fission–fusion dynamics in the spotted hyaena. Anim. Behav. 76, 619–636. (doi:10.1016/j.anbehav.2008.05.001)

20. Holekamp KE, Smale L, Berg R, Cooper SM. 1997 Hunting rates and hunting success in the spotted hyena (Crocuta crocuta). J. Zool. 242, 1–15.

21. Brown AK, Pioon MO, Holekamp KE, Strauss ED. 2021 Infanticide by females is a leading source of juvenile mortality in a large social carnivore. Am. Nat. 198. (doi:10.1086/716636)

22. Lehmann KDS, Montgomery TM, Maclachlan SM, Parker JM, Spagnuolo OS, Vandewetering KJ, Bills PS, Holekamp KE. 2017 Lions, hyenas and mobs (Oh my!). Curr. Zool. 63, 313–322. (doi:10.1093/cz/zow073)

23. Smith JE, Memenis SK, Holekamp KE. 2007 Rank-related partner choice in the fission– fusion society of the spotted hyena (Crocuta crocuta). Behav. Ecol. Sociobiol. 61, 753– 765. (doi:10.1007/s00265-006-0305-y)

24. Strauss ED, Holekamp KE. 2019 Social alliances improve rank and fitness in convention-based societies. Proc. Natl. Acad. Sci. 116, 8919–8924. (doi:10.1073/pnas.1810384116)

25. Gersick AS, Cheney DL, Schneider JM, Seyfarth RM, Holekamp KE. 2015 Long-distance communication facilitates cooperation among wild spotted hyaenas, Crocuta crocuta. Anim. Behav. 103, 107–116. (doi:10.1016/j.anbehav.2015.02.003)

26. Schamberg I, Cheney DL, Clay Z, Hohmann G, Seyfarth RM. 2017 Bonobos use call combinations to facilitate inter-party travel recruitment. Behav. Ecol. Sociobiol. 71. (doi:10.1007/s00265-017-2301-9)

27. Mills MGL. 1990 Kalahari Hyenas: Comparative Behavioral Ecology of Two Species. Caldwell, New Jersey: The Blackburn Press.

28. Holekamp KE, Smale L. 1998 Behavioral development in the spotted hyena. Bioscience 48, 997–1005.

29. Spiegel O, Leu ST, Sih A, Bull CM. 2016 Socially interacting or indifferent neighbours? Randomization of movement paths to tease apart social preference and spatial constraints. Methods Ecol. Evol. 7, 971–979. (doi:10.1111/2041-210X.12553)

30. Holekamp KE, Smith JE, Strelioff CC, Van Horn RC, Watts HE. 2012 Society, demography and genetic structure in the spotted hyena. Mol. Ecol. 21, 613–632.

31. Johnson MP, Tyack PL. 2003 A digital acoustic recording tag for measuring the response of wild marine mammals to sound. IEEE J. Ocean. Eng. 28, 3–12. (doi:10.1109/JOE.2002.808212)

32. Johnson M, De Soto NA, Madsen PT. 2009 Studying the behaviour and sensory ecology of marine mammals using acoustic recording tags: A review. Mar. Ecol. Prog. Ser. 395, 55–73. (doi:10.3354/meps08255)

33. Holekamp KE, Cooper SM, Katona CI, Berry NA, Frank LG, Smale L. 1997 Patterns of association among female spotted hyenas (Crocuta crocuta). J. Mammal. 78, 55–64.

34. Farine DR, Whitehead H. 2015 Constructing, conducting and interpreting animal social network analysis. J. Anim. Ecol. 84, 1144–1163. (doi:10.1111/1365-2656.12418)

35. Holekamp KE, Ogutu JO, Dublin HT, Frank LG, Smale L. 1993 Fission of a spotted hyena clan: consequences of prolonged female absenteeism and causes of female emigration. Ethology 93, 285–299.

36. Smith JE, Van Horn RC, Powning KS, Cole AR, Graham KE, Memenis SK, Holekamp KE. 2010 Evolutionary forces favoring intragroup coalitions among spotted hyenas and other animals. Behav. Ecol. 21, 284–303. (doi:10.1093/beheco/arp181)

37. Ilany A, Holekamp KE, Akçay E. 2021 Rank-dependent social inheritance determines social network structure in a wild mammal population. Science. 352, 348–352. (doi:10.1101/2020.04.10.036087)

38. Papageorgiou D, Christensen C, Gall GEC, Klarevas-Irby JA, Nyaguthii B, Couzin ID, Farine DR. 2019 The multilevel society of a small-brained bird. Curr. Biol. 29, R1120–R1121. (doi:10.1016/j.cub.2019.09.072)

39. Sikes RS, Gannon WL. 2011 Guidelines of the American Society of Mammalogists for the use of wild mammals in research. J. Mammal. 92, 235–253. (doi:10.1644/10-MAMM-F-355.1)

40. Strauss ED, Jensen FH, Gersick AS, Thomas M, Holekamp KE, Strandburg-Peshkin A. 2021 Daily ranging and den usage patterns structure fission-fusion dynamics and social associations in spotted hyenas. Dryad, Dataset. (doi:https://doi.org/10.5061/dryad.0p2ngf22k)

