## Supplementary Materials for "Daily ranging and den usage patterns structure fission-fusion dynamics and social associations in spotted hyenas"

### 1. Supplementary Methods

*Collar specifications*

All collars integrated a Tellus Medium collar (Followit Sweden AB) with a custom-built sound and movement module modified from a DTAG board [29,30], with both modules connected to the main Tellus battery pack (2x LiTh D-cell batteries). The Tellus Medium collar included a VHF transmitter for locating animals in the field, a programmable release system, and a GPS and Iridium satellite module programmed to transmit daily data packages with time, position and battery state of the collar roughly every 3 hours. The sound and movement module incorporated a high-resolution GPS logger (Technosmart Gipsy 5) sampling continuously at 1 Hz, along with a modified DTAG board that recorded continuous single-channel audio (32 kHz), triaxial accelerometer (1000 Hz), and magnetometer (50 Hz) data, and a 64-GB onboard memory module for data storage. A serial connection between the GPS and DTAG board allowed all tags to be synchronized to GPS time, achieving an accuracy of <1s.

*Collar deployment*

To deploy collars, we anesthetized female hyenas with Telazol (6.5 mg/kg) administered in a plastic dart fired from a CO2 powered rifle (Telinject Inc., Saugus, California), then fitted them with collars (Tellus, Sweden). When daily collar updates reported that batteries were running low or stopped responding, a field crew located the collared individual using the VHF transmitter and triggered a remote release system using VHF communication. Collars were subsequently retrieved and their data offloaded in camp.

*GPS data pre-processing*

GPS data were pre-processed to filter out unrealistic locations likely due to GPS error, and to interpolate small gaps in the data. Overall, we took a conservative approach, only removing data points we could be confident were GPS errors and only interpolating very small gaps.

We first converted the latitude and longitude coordinates for each individual to UTM coordinates (eastings and northings) to facilitate further analysis. We then carried out the following steps:

1) *Remove unrealistic speeds:* We removed data that showed unrealistic speeds that were greater than the 99.95 percentile of all observed speeds in the dataset (speeds greater than 14.8 m / s) and replaced the corresponding coordinates with NAs.
2) *Interpolate small gaps:* For any gaps in the data less than 5 seconds in duration, we used linear interpolation to fill them in based on the previous and subsequent location.
3) *Interpolate slightly longer gaps for stationary hyenas:* For gaps in the data greater than 5 seconds but shorter than 5 minutes, if the distance between the previous location of the hyena and its subsequent observed location was less than 5 m, we filled in the missing data with the mean of these two locations.

4) *Remove outlier positions not near other outliers:* We identified positions where either the easting or northing (or both) was below the 0.01% quantile or above the 99.99% quantile for a given individual. If the previous recorded location and the subsequent recorded location were greater than 1 km from these points, we replaced them with NAs.
5) *Remove extreme outliers even if near other outliers:* For any positions outside of a range of the mean + 10 times the standard deviation of each individual's eastings and/or northings, we replaced them with NAs.

In some cases, due to missing data, individuals might not be observed crossing the threshold for determining the start or end of fission-fusion events. These events were included in the total count of events (n = 690 events), but were excluded from all subsequent analyses, for which we only used events where the exact start and end times were known (n = 551 events). In addition, there were some instances of short sequences of missing data within events (one or both GPS devices missed several fixes), which we included as long as the gap was less than 30 minutes and the individuals were still together afterward.

*Accelerometer data pre-processing and computation of VeDBA*

From the raw tri-axial ACC readings (recorded at 1000 Hz), we first downsampled the data to 25 Hz and then computed the vectorial dynamic body acceleration (VeDBA), a proxy for overall activity level similar to the ODBA but that is invariant to rotational transformations (Qasam et al. 2013). For the VeDBA computations we followed Qasem et al 2013, using a window size of 1 second to separate dynamic and static components of acceleration. We then smoothed the VeDBA data with a sliding window of length 1 second and downsampled to 1 measurement per second for the activity synchrony computations.

*Fitting canonical shape of fission-fusion events*

To identify phases of fission-fusion events, we fit a constrained piecewise linear regression constructed of three line segments. These models broke each event into three phases – the *fission phase*, *together phase*, and *fusion phase*. We fit 3 parameters for each event: $b_1$ (the time point at which the *together phase* begins), $b_2$ (the time point at which the *together phase* ends), and $h$ (the average distance between hyenas during the *together phase*).

We imposed constraints to ensure that fits were meaningful and corresponded to the way that we defined fission-fusion events. To match our definition of fission-fusion events, we fixed the starting distance of the fusion phase to be 200m, the slope of the together phase to be 0, and the ending distance of the fission phase to be 200m. To ensure meaningful fits, we constrained h > 0 (average distance between individuals during together phase cannot be negative), $b_1 > t_0$ (*together phase* cannot begin before the start of the entire event), $b_1 < b_2$ (start time of the *together phase* must occur before end time of the *together phase*), and $b_2 < t_f$ (end time of the *together phase* must occur before the end the event).

### 2. Characterizing missing data

GPS tags were programmed to record at a rate of 1 fix per second for 24 hours per day over the entire duration of the study, however there were some gaps in the data. Figure S2 gives a visual representation of the GPS data coverage during the tracking period for each hyena, and Figure S3 shows the percentage of missing GPS data for each hyena across the entire dataset. In addition to some short gaps due to variation in GPS reception, there were also a few instances of 12 hour gaps in GPS recordings due to an integration issue between the audio and GPS tags. Although gaps appeared to occur randomly across days (Figure S4), they occurred more often at specific times of the day, resulting in a non-random pattern in missing data across the day (Figure S5).

Because this daily pattern in missing data showed a similar pattern to our observations of the number of fusion events across the day (Figure 1B), we tested whether this pattern could be an artefact of missing data patterns rather than a true biological result. To do so, we first measured the fraction of pairs of hyenas tracked as a function of hour of the day (Figure S6). We then calculated the total number of seconds hyenas were taking part in fission-fusion events during each hour of the day, and normalized this value by the total number of seconds where both members of the dyad were tracked across all dyads, during each hour of the day. The resulting pattern (Figure S7) looks qualitatively similar to our original result based on the number of fusion events (Figure 1B), with the possible exception that the dip in the middle of the day is not quite as wide.

We also tested whether patterns of missing data across days might somehow bias our permutation test results, which rely on swapping days. For example, if patterns of missing data were correlated among individuals across days, this could lead to an under-representation of fission-fusion events in permuted datasets which would break these correlations. To test whether this could be the case, we computed the total amount of time that both members of a dyad were tracked across our dataset, summing across all dyads. This total number of “dyad-seconds” represents the total amount of time that all dyads were observed in our GPS data and hence could be found to be taking part in fission-fusion events. We then compared this number to the same value computed for 100 permuted reference models. We found that the number of dyad-seconds in the real data fell close to the center of the distribution of number of dyad-seconds across the reference models (Figure S8), suggesting this potential source of bias does not exist in our data. In addition, we found no evidence for autocorrelation in the amount of missing data within individuals from one day to the next (Figure S9). We thus expect that, overall, patterns of missing data should minimally affect our results.

### 3. Supplementary Tables

**Table S1.** Information on collar deployment times for all tracked hyenas.

| Hyena ID | Recording start (GMT + 3) | Recording end (GMT + 3) | Number of days |
|---|---|---|---|
| WRTH | 2017-01-01 00:02:28 | 2017-02-08 12:57:57 | 38.5 |
| BORA | 2017-01-01 00:02:04 | 2017-02-07 17:54:31 | 37.7 |
| BYTE | 2017-01-01 00:02:34 | 2017-02-09 18:29:50 | 39.8 |
| MGTA | 2017-01-01 00:00:34 | 2017-02-04 13:27:08 | 34.6 |
| FAY | 2017-01-01 00:01:22 | 2017-02-10 23:00:32 | 41.0 |

**Table S2.** Metrics used to characterize fission-fusion events

| Metric | Description |
|---|---|
| Duration | **Elapsed time from the beginning to the end of the event.** The beginning of the event is defined as the time at which the distance between the two individuals falls below 200 m, and the end as the time at which this distance increases above 200 m. |
| Displacement | **Total displacement of the individuals during the together phase**. To produce a single metric, each individual's displacement is calculated separately, and the smallest of these two values was used. |
| Directional synchrony | **Correlation between the headings of the two individuals during the together phase.** The heading of each individual at every moment in time was calculated as the unit vector pointing from its current position to the position it occupied after moving a distance of 5 m (this spatial threshold was used to reduce the influence of GPS jitter). The directional synchrony was then calculated by taking the dot product between the headings of the two individuals, and averaging this value across all time points during the together phase. To reduce the influence of times when one or more hyena was stationary, we excluded from our calculations times when either individual failed to move at least 5 meters during the subsequent 10 seconds. |
| Activity synchrony | **Correlation between the vectorial dynamic body association (veDBA) of the two individuals during the together phase.** The veDBA is a measure of overall activity level of an individual. |
| Distance from den at start | **Distance between hyenas and the closest den at the beginning of the event.** Here, the smaller distance from the den of the two hyenas was used. |
| Distance from | **Distance between the hyenas and the closest den at the end of the** |

| | |
|---|---|
| den at end | **event.** Here, the smaller distance from the den of the two hyenas was used. |

### 4. Supplementary Figures

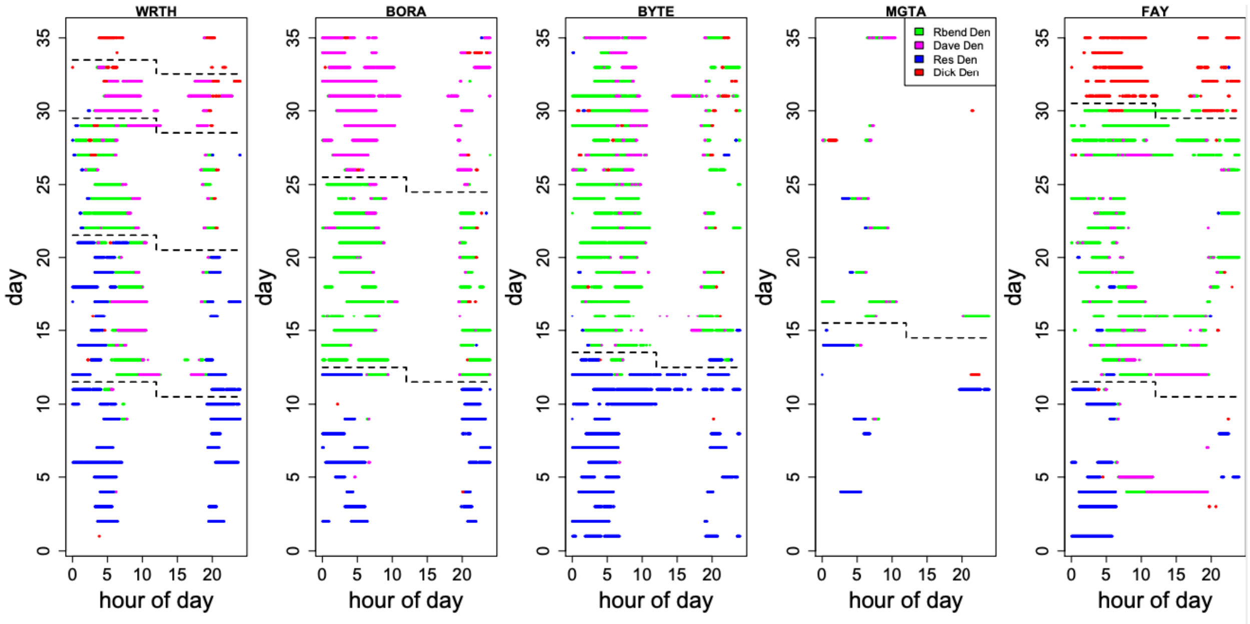

**Figure S1. Den usage by the five tracked hyenas over the course of the study period.** Colored dots represent when each hyena (different panels) was within 200 m of a given den (colors - see legend in fourth panel from left) as a function of day within the study period (y axis) and time of day (x axis). Dashed lines indicate block of time when each hyena was more or less consistent in its den usage - in reference models, days were only swapped within these blocks. Note that blocks started at noon on a given day (represented by the zig-zag shape of the dotted lines).

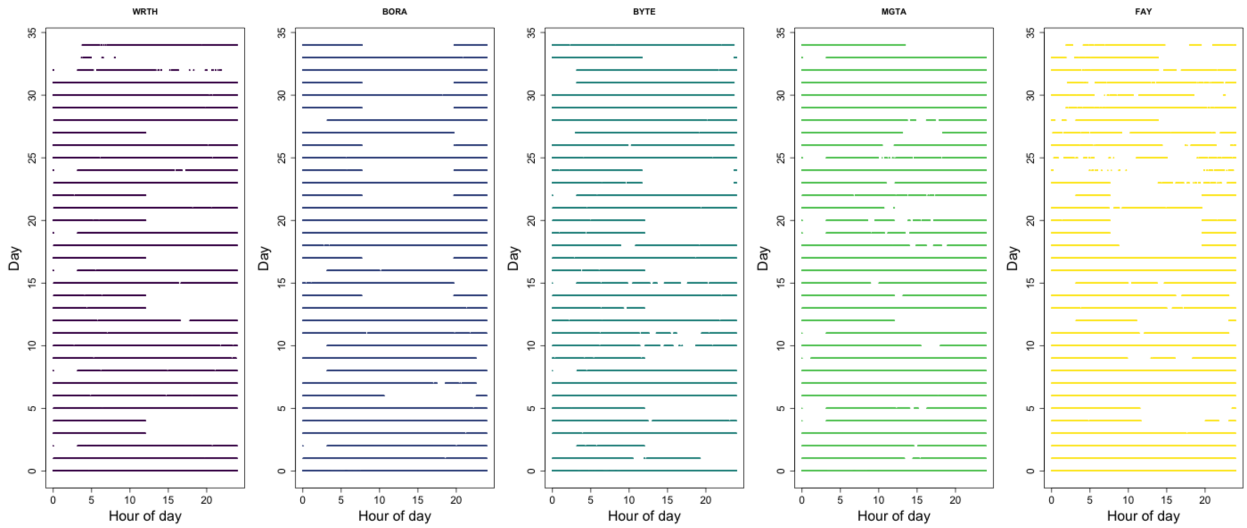

**Figure S2. Missing data for each individual throughout the study period.** Colored bars represent times when the hyena was tracked on each day (y-axis) at each hour of the day (x-axis), with empty regions showing instances of missing data.

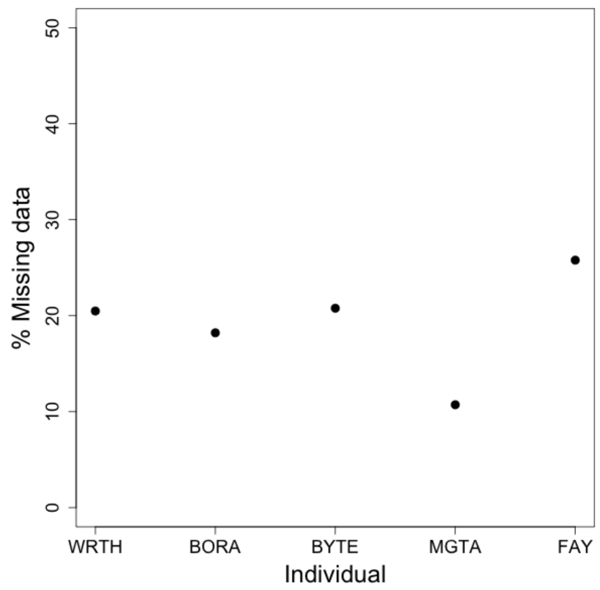

**Figure S3. Total percentage of missing data for each individual studied.**

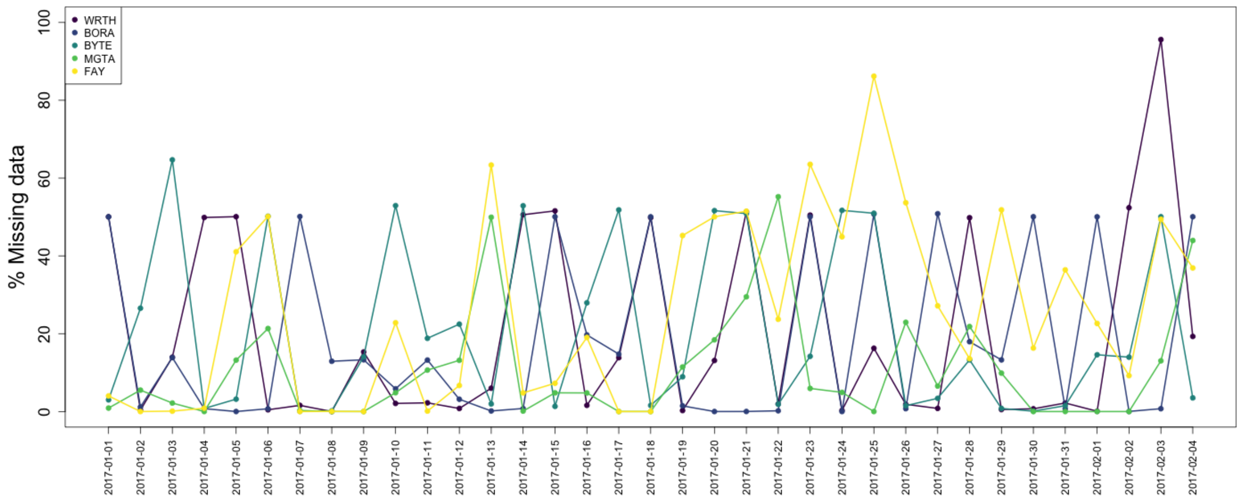

**Figure S4. Total amount of missing data for each individual (colored lines) on each day (x-axis).**

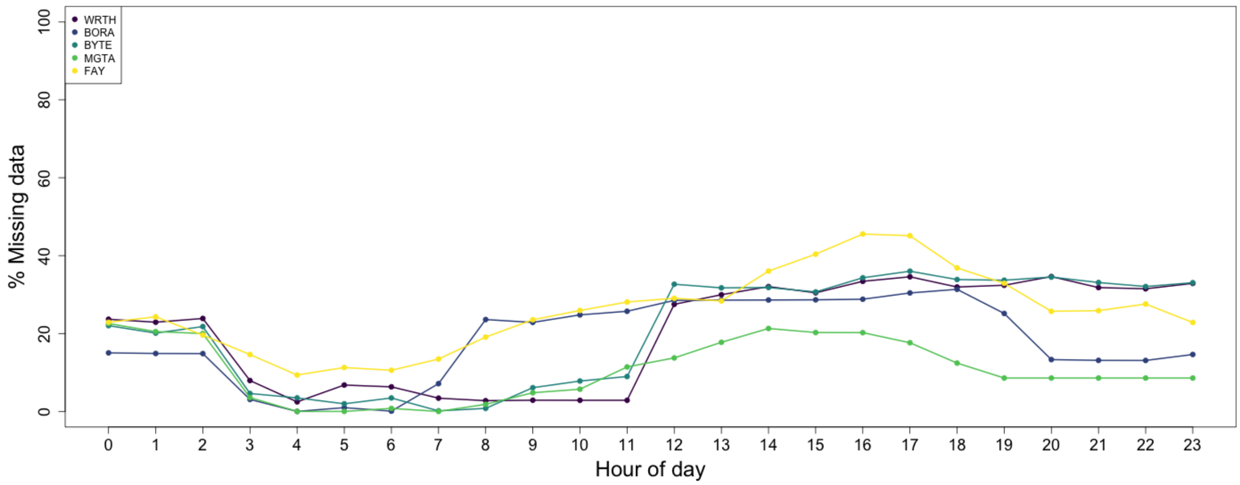

**Figure S5. Percentage of missing data (y-axis) for each hour of the day (x-axis), for each individual (colored lines).** Missing data were more common at midday - evening.

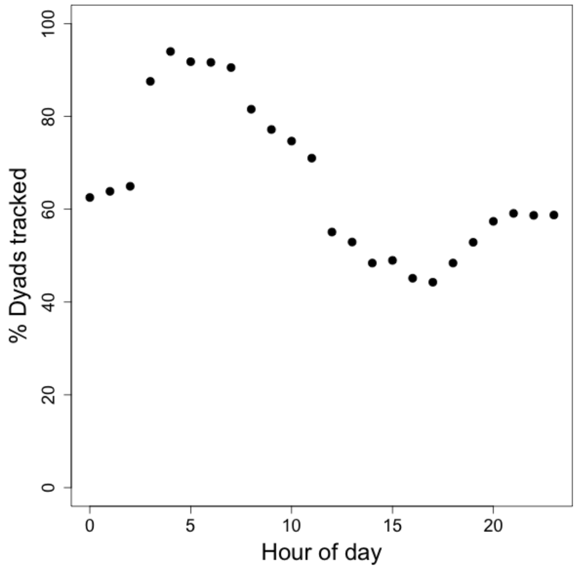

**Figure S6. Percentage of dyads that were tracked for each hour of the day, across the entire study period.**

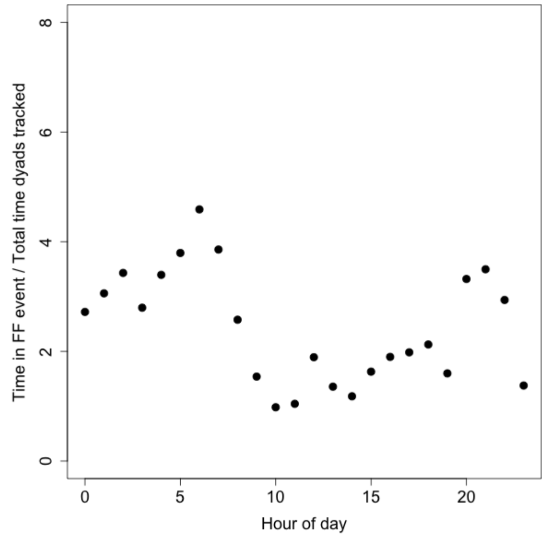

**Figure S7. Total percentage of time dyads were engaged in fission-fusion events as a function of time of day.**

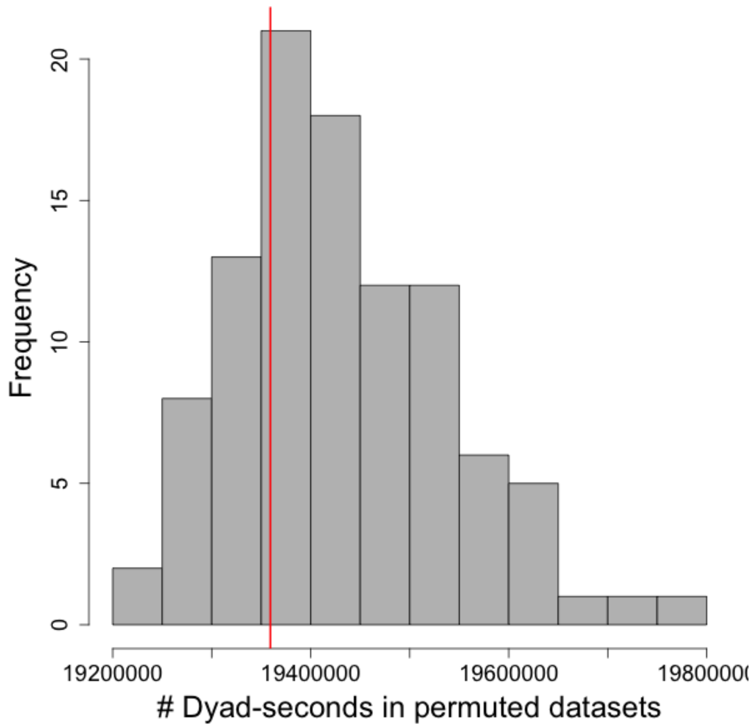

**Figure S8. Total amount of time dyads were tracked in the permuted reference models (histogram) compared to the real data (red line).**

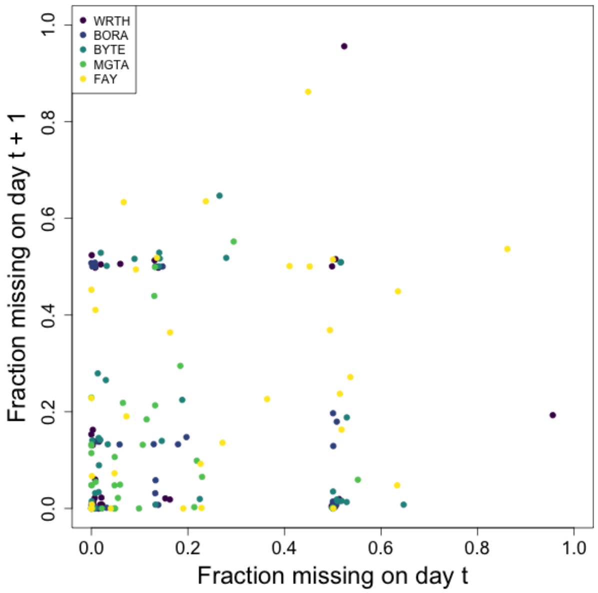

**Figure S9. Lack of temporal autocorrelation across days in the amount of missing data.** Plot shows fraction of missing data on a given day *t* on the x-axis and the fraction of missing data on the subsequent day *t* + 1 on the y-axis. Colors represent different individuals.

### 5. Alternative Parameterizations

Here we provide results from analyses with alternative parameterizations. These plots correspond to the Figures 1 – 5 in the main text. The parameters we varied here are the distance thresholds used for identifying fission-fusion events. The **inner threshold** sets the criteria for identification of a fission-fusion event, and the **outer threshold** determines when the event starts and stops. A fission-fusion event occurs when the distance between two individuals crosses the inner threshold. The event starts when the distance between the two individuals first crosses the outer threshold and ends when the distance between the two individuals again crosses the outer threshold after crossing the inner threshold. In the main analysis, these (inner / outer) thresholds were 100m / 200m. Here we present results for two alternative parameterizations, 50m / 100m and 200m / 300m.

*50m / 100m*

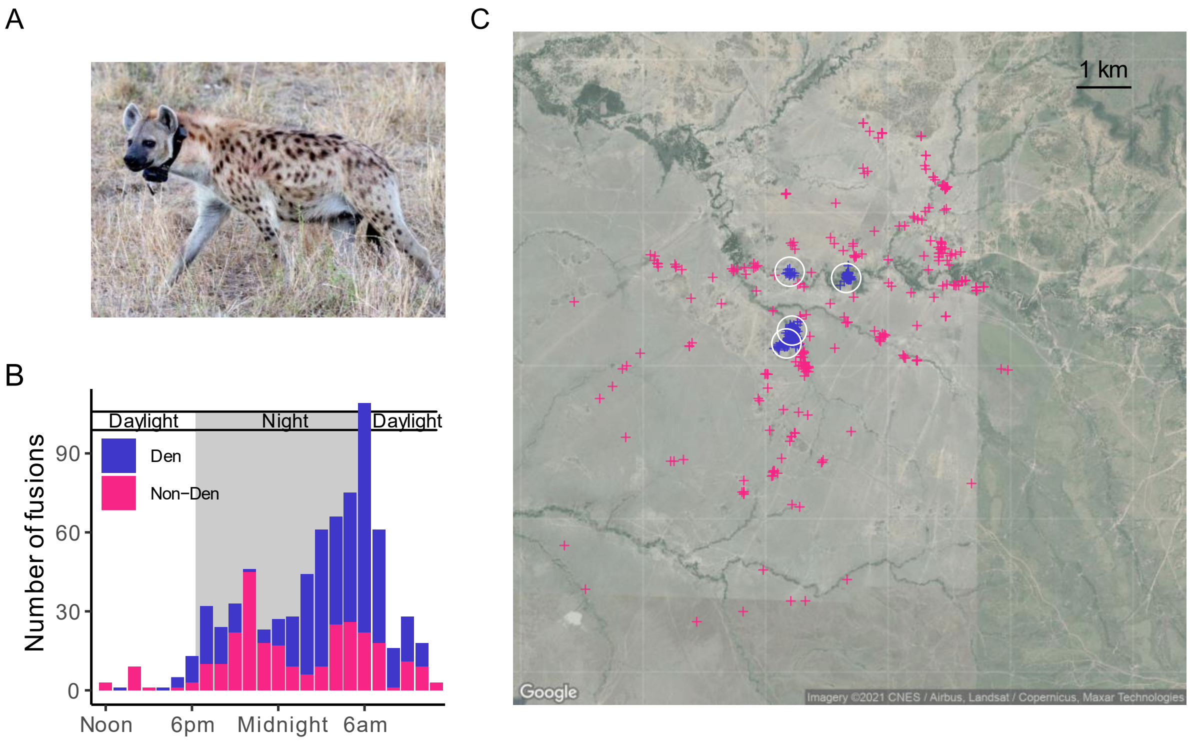

**Figure S10. Spatial and temporal patterns of fission-fusion in spotted hyenas.** (A) Female hyena wearing a tracking collar. (B) Time of day and (C) locations of the starts of fission-fusion events across all hyena pairs. Color specifies whether events started a den (blue) or not (magenta). White circles represent locations of the four communal dens in use during the study period. See also Supplementary Video 1.

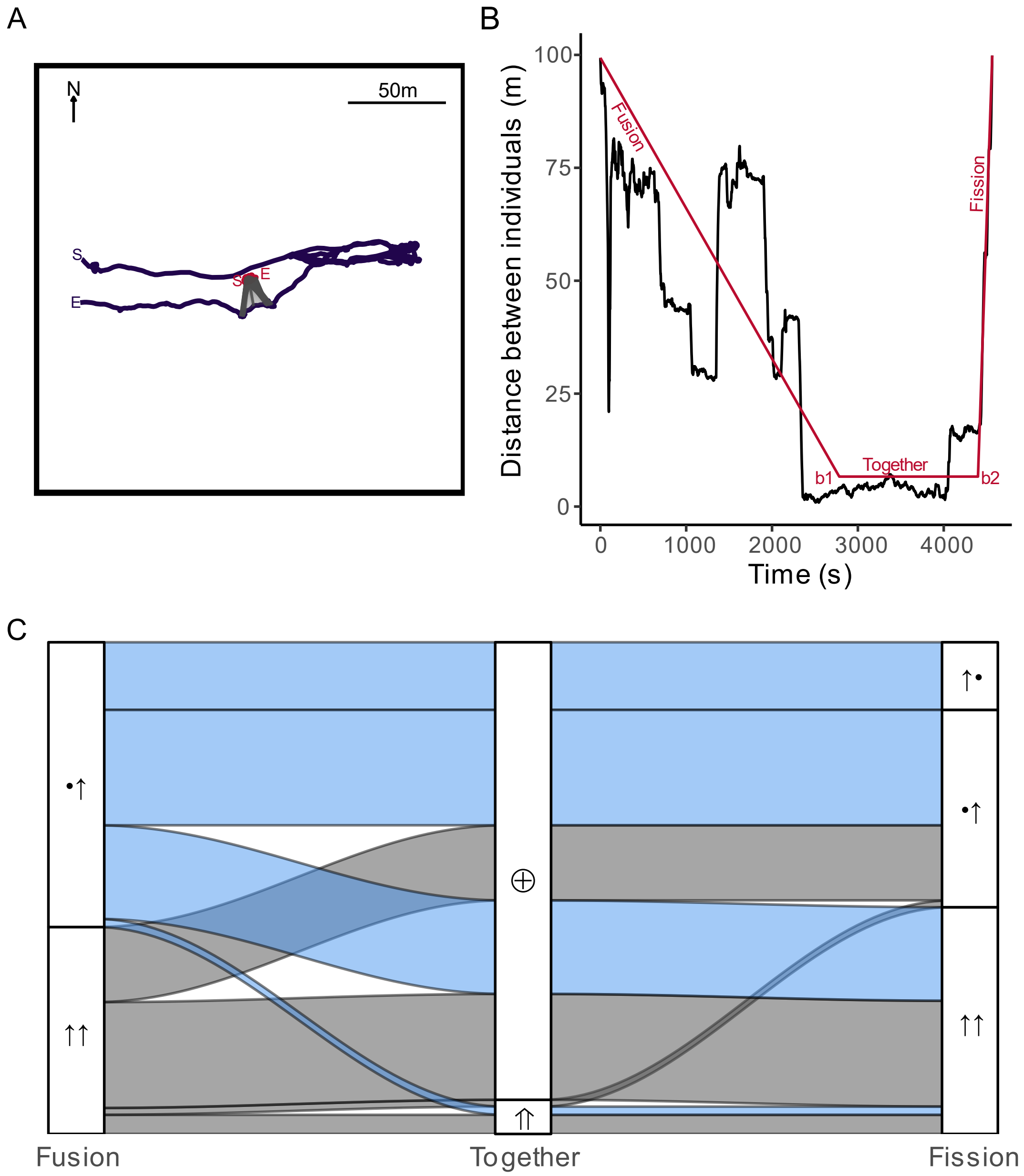

**Figure S11**. **Example fission-fusion event and frequencies of event types.** (A) Example trajectories of two individuals during an extracted fission-fusion event. Gray lines connect time-matched GPS points during the *together phase*. See Supplementary Videos for animated examples. (B) Distance between the two individuals over time (black) and fitted piecewise regression model (red). Break points are used to identify the three phases for each event. (C) Alluvial plot of the frequencies of transition motifs between different categories of the three phases. Symbols indicate the movement patterns of the two individuals involved in the event (• = stationary, ↑ = moving, ⊕ = local, ⇑ = traveling). Note that asymmetrical fissions can occur in two ways: either the two individuals show the same movement patterns as in the fusion phase (•↑), or the individuals reverse movement patterns (↑•).

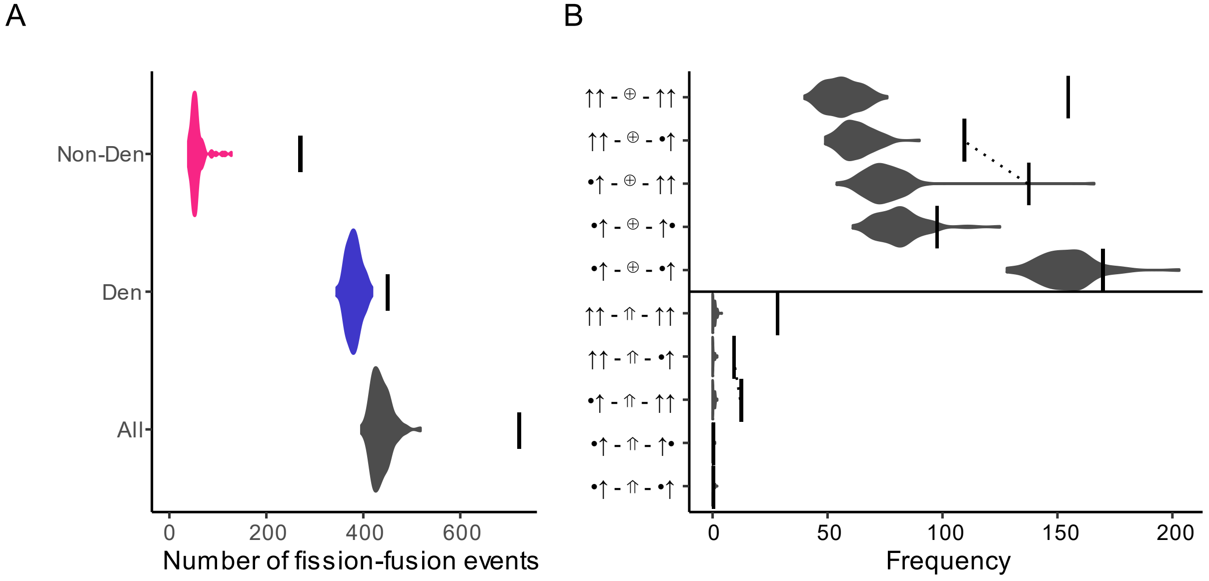

**Figure S12. Comparison of the number of fission-fusion events in real data vs. reference models preserving den attendance and daily ranging patterns.** (A) Overall number of events observed across real data (vertical lines) and reference models (violin plots) across all events, den events, and non-den events. (B) Frequency of events (x-axis) broken down by type (y-axis) in the real data (vertical lines) as compared to the reference models (violin plots). Y-axis labels represent the behavior of the two individuals during the three phases (from left to right: fusion, together, fission) for each event type. Dotted lines connect pairs of event types that are essentially time-reversed versions of each other, to highlight the asymmetry between fusions and fissions (see text).

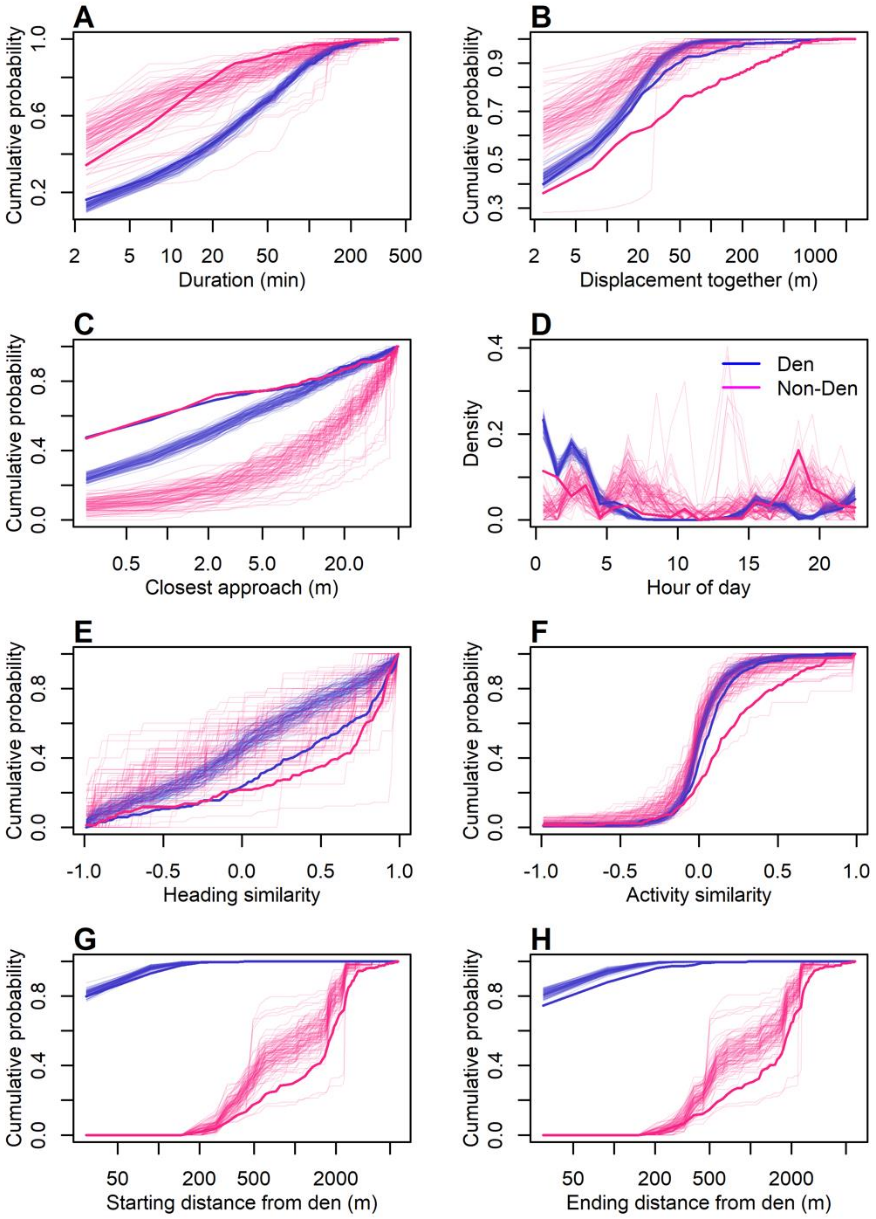

**Figure S13. Reference models reproduce some but not all properties of observed fission-fusion events.** Comparison of the detailed properties of fission-fusion events in the real data (thick lines) and in reference models (thin lines), broken up by whether the event occurred in the vicinity of a den (blue) or not (magenta). Plots show the cumulative distribution of each metric (x-axis labels) across all events. Note that panel D shows the distribution rather than the cumulative distribution.

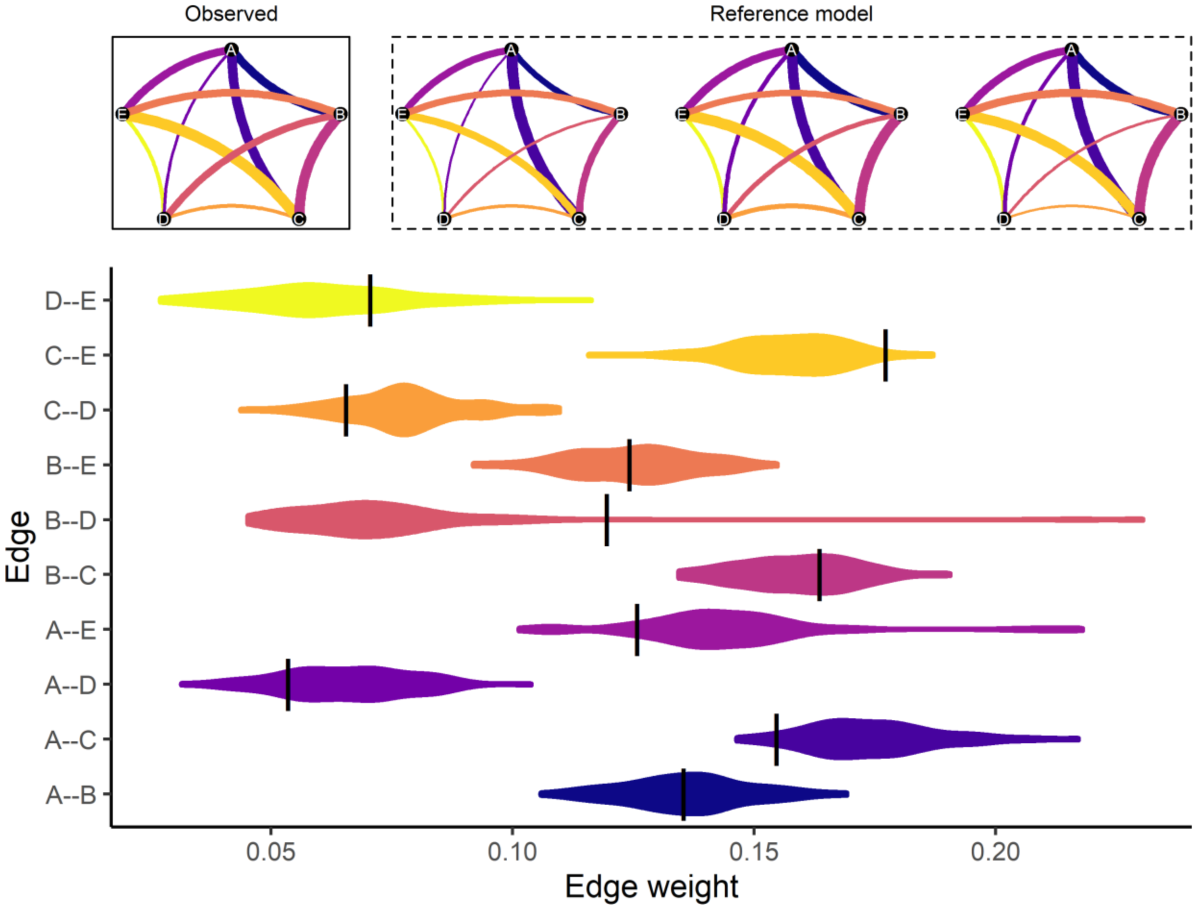

**Figure S14. Reference models reproduce differentiated relationships found in observed social networks built from fission-fusion events.** Black lines indicated observed edge weight representing frequency of association. Violins depict distributions of edge weights in 100 instances of the reference model.

*200m / 300m*

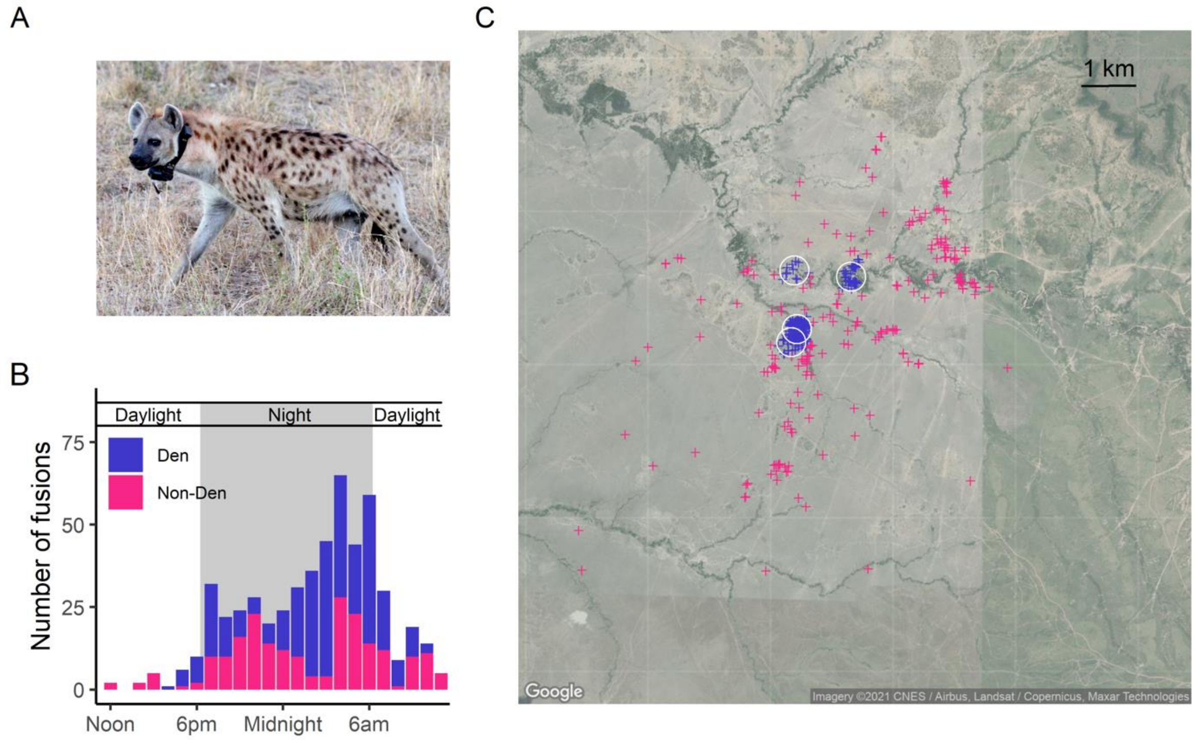

**Figure S15. Spatial and temporal patterns of fission-fusion in spotted hyenas.** (A) Female hyena wearing a tracking collar. (B) Time of day and (C) locations of the starts of fission-fusion events across all hyena pairs. Color specifies whether events started a den (blue) or not (magenta). White circles represent locations of the four communal dens in use during the study period. See also Supplementary Video 1.

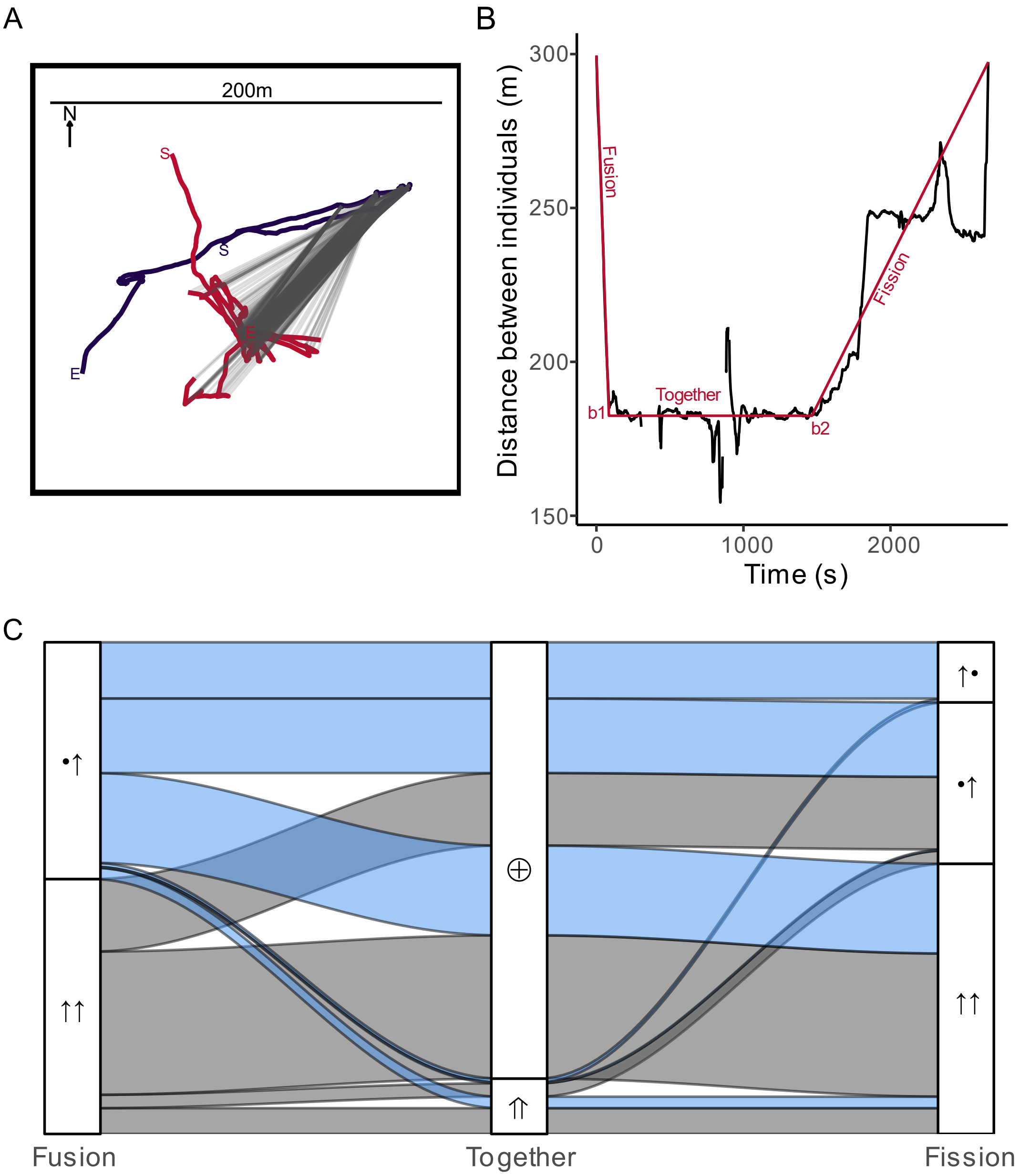

**Figure S16**. **Example fission-fusion event and frequencies of event types.** (A) Example trajectories of two individuals during an extracted fission-fusion event. Gray lines connect time-matched GPS points during the *together phase*. See Supplementary Videos for animated examples. (B) Distance between the two individuals over time (black) and fitted piecewise regression model (red). Break points are used to identify the three phases for each event. (C) Alluvial plot of the frequencies of transition motifs between different categories of the three phases. Symbols indicate the movement patterns of the two individuals involved in the event (• = stationary, ↑ = moving, ⨁ = local, ⇑ = traveling). Note that asymmetrical fissions can occur in two ways: either the two individuals show the same movement patterns as in the fusion phase (•↑), or the individuals reverse movement patterns (↑•).

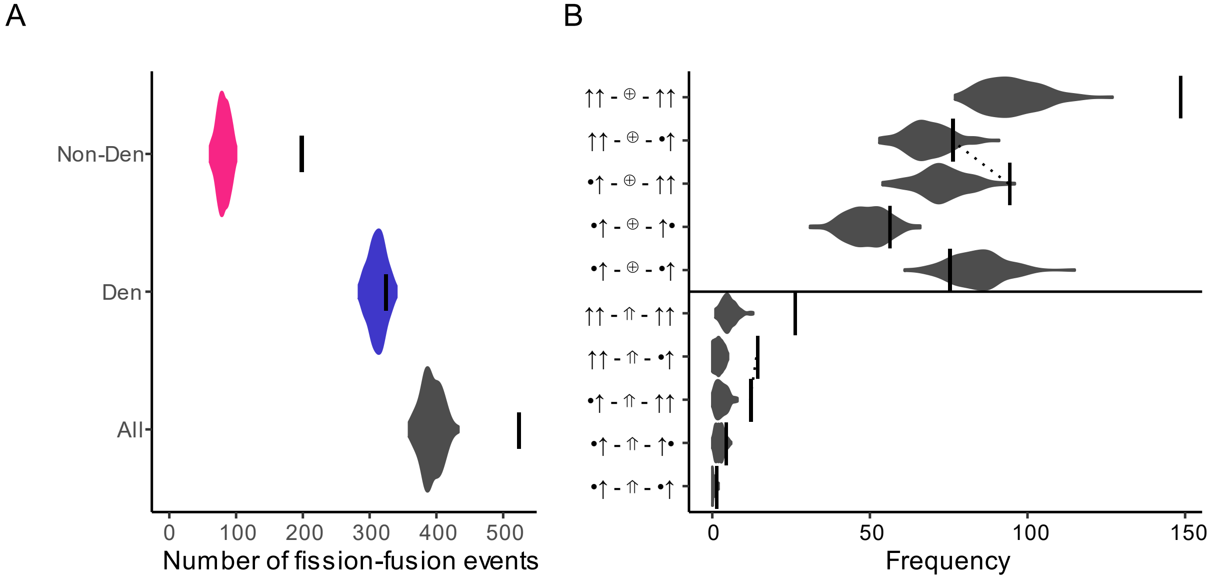

**Figure S17. Comparison of the number of fission-fusion events in real data vs. reference models preserving den attendance and daily ranging patterns.** (A) Overall number of events observed across real data (vertical lines) and reference models (violin plots) across all events, den events, and non-den events. (B) Frequency of events (x-axis) broken down by type (y-axis) in the real data (vertical lines) as compared to the reference models (violin plots). Y-axis labels represent the behavior of the two individuals during the three phases (from left to right: fusion, together, fission) for each event type. Dotted lines connect pairs of event types that are essentially time-reversed versions of each other, to highlight the asymmetry between fusions and fissions (see text).

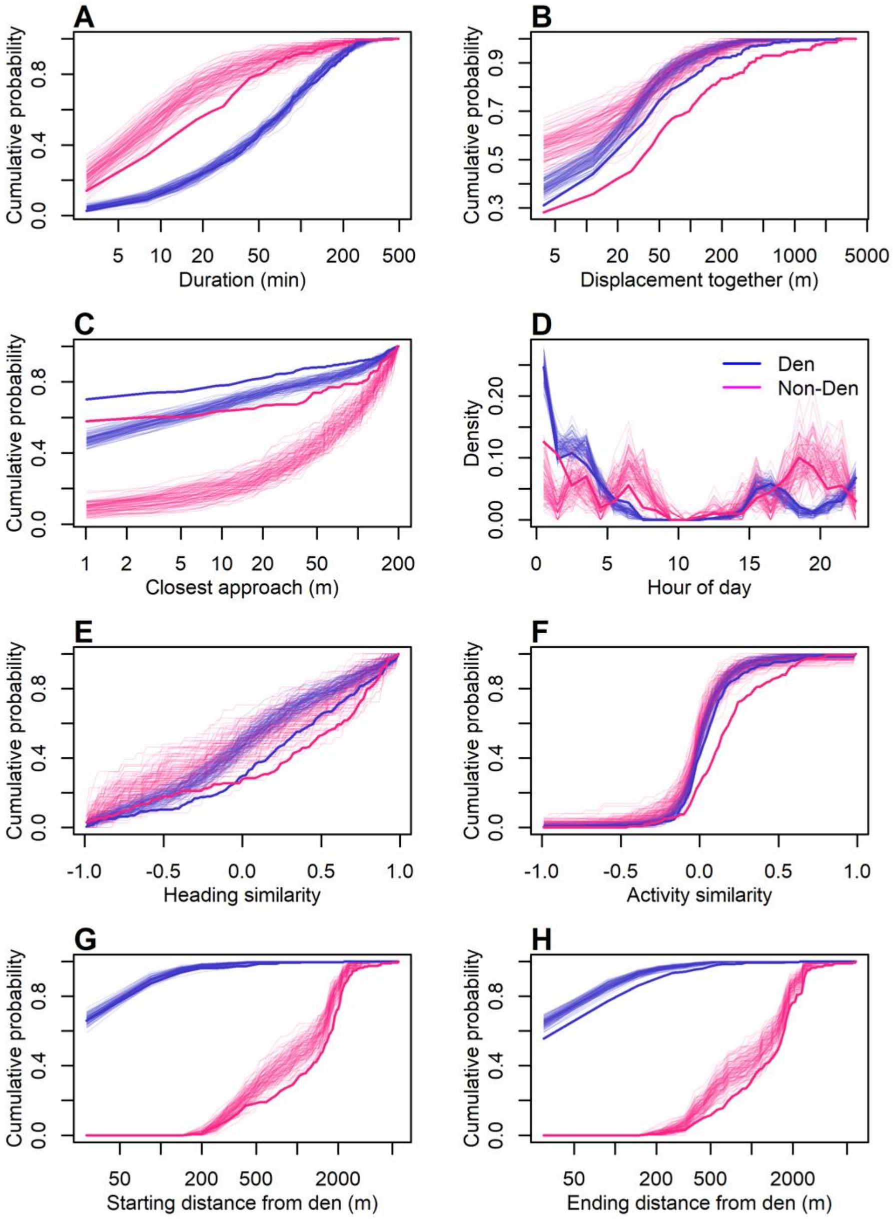

**Figure S18. Reference models reproduce some but not all properties of observed fission-fusion events.** Comparison of the detailed properties of fission-fusion events in the real data (thick lines) and in reference models (thin lines), broken up by whether the event occurred in the vicinity of a den (blue) or not (magenta). Plots show the cumulative distribution of each metric (x-axis labels) across all events. Note that panel D shows the distribution rather than the cumulative distribution.

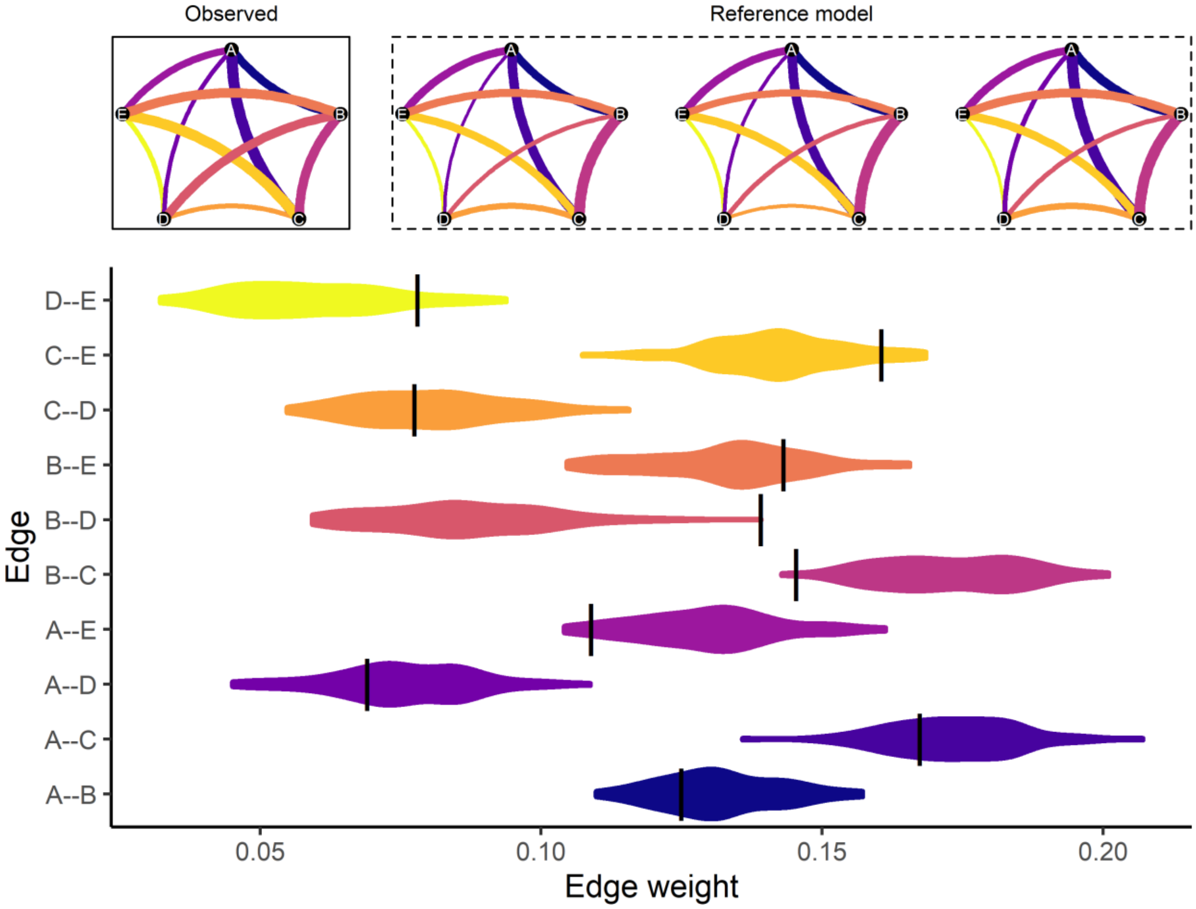

**Figure S19. Reference models reproduce differentiated relationships found in observed social networks built from fission-fusion events.** Black lines indicated observed edge weight representing frequency of association. Violins depict distributions of edge weights in 100 instances of the reference model.

### 6. Supplementary Videos

**Supplementary Video 1.**
Movement of the five study individuals over the entire duration of the study. White circles mark the locations of four communal dens that were in use at some point during the study (see Supplementary Figure S1).

**Supplementary Video 2**.
Example fission-fusion event of type: •↑ - ⊕ - •↑. The *together phase* is indicated by the darker points and the trails behind each individual.

**Supplementary Video 3**.
Example fission-fusion event of type: •↑ - ⇑ - ↑↑. The *together phase* is indicated by the darker points and the trails behind each individual.

**Supplementary Video 4**.
Example fission-fusion event of type: ↑↑ - ⨁ - ↑↑. The *together phase* is indicated by the darker points and the trails behind each individual.
